# DNA elements tether canonical Polycomb Repressive Complex 1 to human genes

**DOI:** 10.1101/2023.01.12.523763

**Authors:** Juan I. Barrasa, Tatyana G. Kahn, Moa J. Lundkvist, Yuri B. Schwartz

## Abstract

Development of multicellular animals requires epigenetic repression by Polycomb group proteins. The latter assemble in multi-subunit complexes, of which two kinds, Polycomb Repressive Complex 1 (PRC1) and Polycomb Repressive Complex 2 (PRC2), act together to effect the repression of key developmental genes. How PRC1 and PRC2 recognize specific genes remains an open question. Here we report systematic identification of DNA elements that tether canonical PRC1 to human developmental genes. Their analysis indicates that sequence features associated with PRC1 tethering differ from those that favour PRC2 binding. Throughout the genome, the two kinds of sequence features mix in different proportions to yield a gamut of DNA elements that range from those tethering predominantly PRC1 to ones capable of tethering both PRC1 and PRC2. The emerging picture is similar to paradigmatic targeting of Polycomb complexes by Polycomb Response Elements (PREs) of *Drosophila* but providing for greater plasticity.

## INTRODUCTION

Multicellular animals rely on epigenetic mechanisms to repress alternative gene expression programs when their cells acquire specialized functions. This is common during embryonic development and critical in later life to replenish specific cell pools from multipotent stem cells. Polycomb group proteins make up the epigenetic system most widely used to represses alternative gene expression programs in differentiated cells (Piunti and Shilatifard, 2021; Schuettengruber et al., 2017).

First discovered in fruit flies *Drosophila melanogaster* as critical regulator of homeotic selector genes, the Polycomb system was later shown to repress hundreds of genes encoding transcriptional regulators, morphogenes and signalling molecules that are involved in all main developmental pathways (Boyer et al., 2006; Lee et al., 2006; Schwartz et al., 2006). Many of the same genes are targeted by the system in *Drosophila* and mammalian cells, which suggests that Polycomb group proteins were co-opted to regulate these genes long time ago, before the insect and vertebrate lineages split.

Early biochemical studies have shown that Polycomb group proteins assemble in two kinds of evolutionarily conserved Polycomb Repressive Complexes, PRC1 and PRC2 (Czermin et al., 2002; Kuzmichev et al., 2002; Levine et al., 2002; Muller et al., 2002; Shao et al., 1999). Mammalian version of PRC1 consists of a heterodimer between RING2 (or its paralogue RING1) and MEL18 or closely related BMI1; a chromodomain-containing subunit represented by either CBX2, CBX4, CBX6, CBX7 or CBX8; one of the Polyhomeotic-like proteins (PHC1, PHC2 or PHC3) and SCMH1 (or its paralogues SCML1, SCML2).

PRC2 complexes contain a core of four subunits: EZH2 (or closely related EZH1), SUZ12, EED and RBBP7 (or the related protein RBBP4). The PRC2 core may further associate with alternative sets of auxiliary subunits: PHF1 (or its paralogues PHF19 and MTF) (Alekseyenko et al., 2014; Chen et al., 2018; Conway et al., 2018; Nekrasov et al., 2007) or JARID2 and AEBP2 (Li et al., 2010; Pasini et al., 2010; Peng et al., 2009; Shen et al., 2009). All PRC2 variants can methylate histone H3 at Lysine 27 (H3K27) and tri-methylation of H3K27 is essential for PRC2 contribution to epigenetic repression by Polycomb system (Pengelly et al., 2013; Sankar et al., 2022).

In addition to the “canonical” complexes described above, the RING1 or RING2 subunits of PRC1 are incorporated in several other complexes sometimes referred to as “non-canonical” or “variant” PRC1 (Gao et al., 2012; Hauri et al., 2016; Kloet et al., 2016; Tavares et al., 2012). Along with RING1 or RING2, these complexes contain RYBP (or closely related YAF2) protein but lack chromodomain containing (CBX) and Polyhomeotic-like (PHC) subunits. These complexes may include MEL18 or BMI1 but, predominantly, incorporate one of the closely related PCGF1, PCGF3, PCGF5 or PCGF6 proteins instead. Several constellations of additional subunits distinguish various RING-RYBP complexes from each other. Once thought to be a vertebrate-specific novelty, the orthologous RING-RYBP complexes were recently identified in *Drosophila* (Kang et al., 2022) suggesting that both canonical PRC1 and RING-RYBP complexes are evolutionarily old (Gahan et al., 2020). PRC1 and RING-RYBP complexes can monoubiquitylate histone H2A at Lysine 119 although the former appears less active in reactions reconstituted *in vitro* (Gao et al., 2012; Rose et al., 2016; Taherbhoy et al., 2015). Multiple lines of evidence argue that canonical PRC1 is critical for repression of developmental genes (Akasaka et al., 2001; Bel et al., 1998; Isono et al., 2005; Jürgens, 1985; Oktaba et al., 2008; van der Lugt et al., 1994). To what extent the monoubiquitylation of H2A or various RING-RYBP complexes contribute to the repression is a subject of debate (Blackledge et al., 2020; Bonnet et al., 2022; Fursova et al., 2019; Illingworth et al., 2015; Lee et al., 2015; Pengelly et al., 2015; Scelfo et al., 2019; Tamburri et al., 2020).

Developmental genes repressed by Polycomb mechanisms are bound by PRC1 and PRC2 and enriched in tri-methylated H3K27 (H3K27me3). What molecular mechanisms enable specific binding of the complexes to these genes is an important open question. Neither canonical PRC1 nor PRC2 have subunits that contain sequence specific DNA binding domains. In *Drosophila*, genes regulated by Polycomb system contain specialized Polycomb Response Elements (PREs). These are short (∼1kb) DNA elements, which can be pinpointed from genomic Chromatin Immunoprecipitation (ChIP) profiles as distinct peaks co-occupied by PRC1 and PRC2 (Schwartz et al., 2006; Schwartz et al., 2010). PREs are sufficient to generate new binding sites for PRC1 and PRC2 when integrated elsewhere in the fly genome. Conversely, their deletions cause stochastic re-activation of associated genes in cells where those are normally inactive or transcribed at low levels (Ogiyama et al., 2018; Sipos et al., 2007). Which DNA elements direct PRC1 and PRC2 to specific genes in mammals is less clear. Most of what we know about genomic distribution of mammalian PRC1 and PRC2 comes from studies of mouse embryonic stem cells. In these cells, PRC1 and PRC2 seem to bind repressed genes in broad and virtually identical fashion with no features to distinguish potential tethering DNA elements (Fursova et al., 2019; Healy et al., 2019; Tamburri et al., 2020). Multiple observations indicate that PRC2 prefers to bind at genomic sites enriched in stretches of unmethylated CpG di-nucleotides (Jermann et al., 2014; Lynch et al., 2012; Mendenhall et al., 2010). PCL, JARID2 and AEBP2 subunits of PRC2 have micromolar affinity to DNA, with possible preference for CpG (Choi et al., 2017; Li et al., 2010; Li et al., 2017; Wang et al., 2017), which may contribute to this binding bias (Healy et al., 2019; Hojfeldt et al., 2019). However, only a fraction of genomic unmethylated CpG-rich sites is bound by PRC2, which argues that other determinants must contribute to the binding specificity.

A handful of DNA elements capable of tethering PRC1 were serendipitously identified within mouse and human genes (Cameron et al., 2018; Sing et al., 2009; Woo et al., 2010). Of those, 1kb-long PRC1 Tethering Element (PTE) upstream of the human *Cyclin D2* (*CCND2*) gene is particularly interesting (Cameron et al., 2018). It was pinpointed in NT2-D1 embryonic teratocarcinoma cells as a strong localized peak of ChIP enrichment with antibodies against PRC1 subunits but disproportionally weak ChIP signals for PRC2 and H3K27me3. Processes linked to high transcriptional activity of *CCND2* in the NT2-D1 cells interfere with PRC2 binding to the locus but do not affect PRC1 tethering by the PTE. Consistently, in TIG-3 human embryonic fibroblast cells where *CCND2* is repressed by Polycomb mechanisms, ChIP-signals for PRC2 and H3K27me3 are abundantly present at a CpG-island adjacent to the *CCND2* PTE (Cameron et al., 2018). In these cells, PRC1 is also present at the CpG-island. However, the corresponding ChIP signals are much weaker than those at the PTE. Transgenic experiments showed that DNA underneath *CCND2* PTE was sufficient to re-create strong PRC1 binding when integrated elsewhere in the genome. PRC2 was also detectable at the transgenic PTE. However, in contrast to PRC1, ChIP signals for PRC2 and associated H3K27me3 were low and just above the genomic background. Taken together, these observations argue that the efficient binding of PRC1 and PRC2 to *CCND2*relies on distinct DNA features and that, at this locus, separable DNA elements combine their inputs to achieve coordinated presence of the two complexes.

How many other developmental genes have PTEs? Are these elements important for epigenetic repression of the cognate genes? What sequence features promote PRC1 binding? To address these questions we performed an unbiased genomic screen, which uncovered PTEs at hundreds of genes encoding regulators of organ development, cell fate commitment and pattern specification. We found that PTEs repress transcription of associated genes and that DNA sequence features required to tether PRC1 differ from those that drive PRC2 binding.

## RESULTS

We identified *CCND2* PTE by genomic mapping of Polycomb group proteins on chromosomes 8, 11 and 12 of human NT2-D1 cells as a site with disproportionally strong ChIP signal for PRC1 but nearly background signals for PRC2 and H3K27me3 (Cameron et al., 2018). The PTE stood out because, in these cells, some processes, linked to the *CCND2* transcription, interfered with PRC2 binding to the locus but did not affect PRC1 tethering by the PTE (Cameron et al., 2018). We reasoned that the same approach might reveal additional PTEs if expanded genome wide. To this effect, we mapped MEL18 (core PRC1 subunit) and SUZ12 (core PRC2 subunit) binding along the genomes of NT2-D1 and TIG-3 cells using Chromatin Immunoprecipitation coupled to massively parallel sequencing of precipitated DNA (ChIP-seq). We performed two independent ChIP-seq experiments for each cell line and antibody and sequenced the DNA from corresponding chromatin input materials to control for potential sample processing biases.

In preliminary experiments, we noticed that size-selection of immunoprecipitated DNA fragments, commonly used during ChIP-sequencing library preparation, distorts immunoprecipitation profiles compared to those derived from the same ChIP reaction by qPCR. We, therefore, omitted the size-selection step and, instead, fragmented immunoprecipitated DNA enzymatically to 180bp prior to ligation of adapters for Illumina sequencing (see Materials and Methods for details).

We then used the ChIP-seq profiles to search the genome of NT2-D1 cells for sites that resemble *CCND2*. That is the sites that: *i*) displayed strong MEL18 ChIP-seq signal (>3000 RPKM) comparable to that at the *CCND2* locus, *ii*) had the ratio between MEL18 and SUZ12 ChIP-seq signals higher than that at *CCND2*, *iii*) showed strong binding of both MEL18 and SUZ12 (both > 2000 RPKM) in TIG-3 cells. Three genomic sites, located in the vicinity of *ZIC2*, *NR6A1* and *SOX21*genes, matched this criteria (Figure 1A-B). Of those, the highest MEL18/SUZ12 ChIP-seq signal ratio corresponded to the site downstream of the *ZIC2* gene. *ZIC2* encodes a human homolog of the *Drosophila* Odd-paired (Opa) transcription factor (Brown et al., 1998) and is disrupted in ∼5% of holoprosencephaly patients (Dubourg et al., 2018).

**Figure 1.**
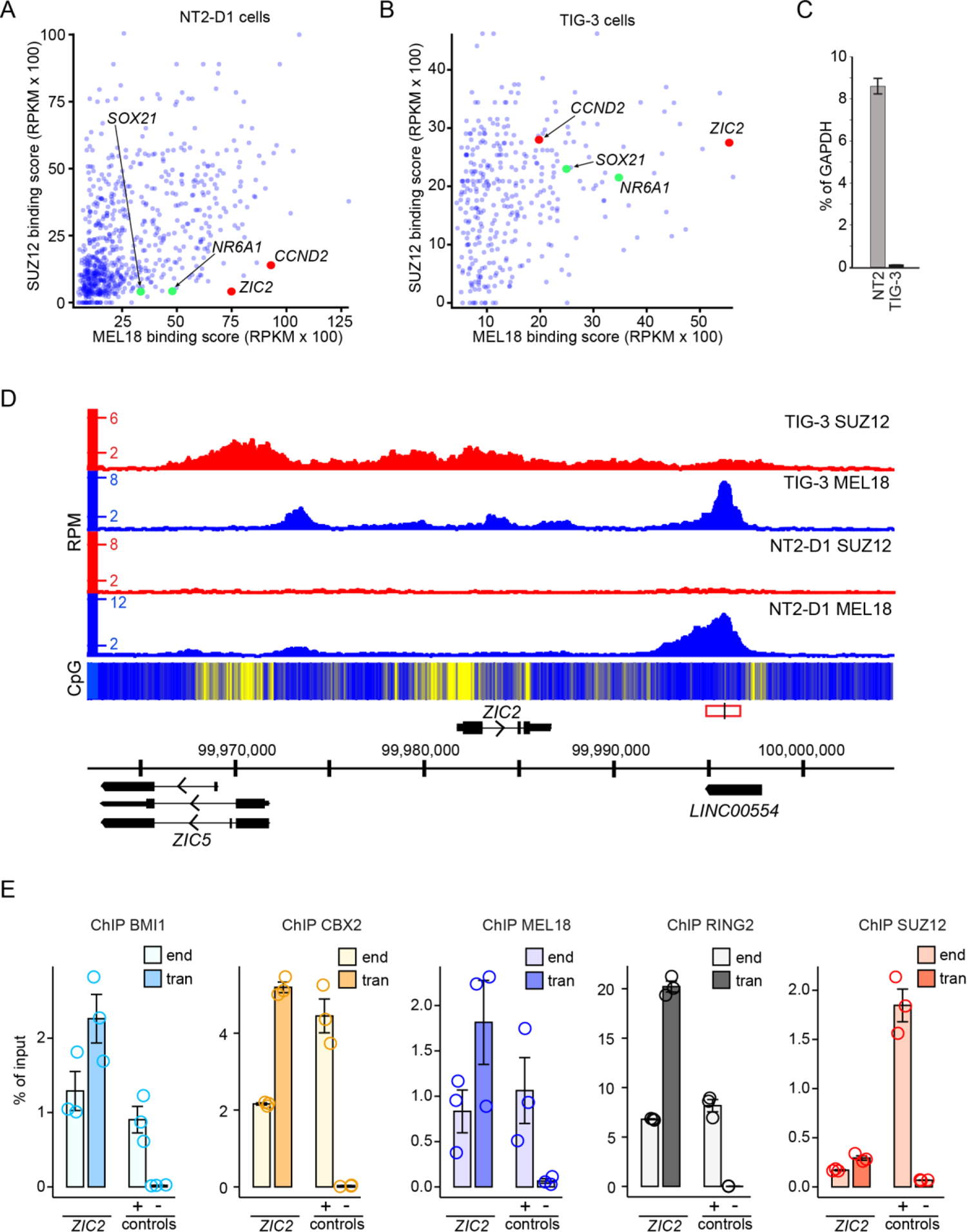
*ZIC2* locus contains a PRC1 tethering element. MEL18 and SUZ12 binding scores for NT2-D1 (**A**) and TIG-3 (**B**) cells at regions significantly enriched by immunoprecipitation of NT2-D1 chromatin with anti-MEL18 antibodies. Each point of the scatter-plot represents unique region. Regions encompassing *CCND2* (Cameron et al., 2018) and *ZIC2* PTEs are marked in red. *SOX21* and *NR6A1* regions, which also matched our screening criteria, are marked in green. **C.** RT-qPCR measurements show that *ZIC2* gene is transcriptionally active in NT2-D1 (NT2) cells but inactive in TIG-3 cells. The bar-plot displays average cDNA counts normalized to the transcription of the housekeeping *GAPDH* gene. Whiskers indicate the scatter between two independent experiments. **D.** Screen-shot of the MEL18 and SUZ12 ChIP-seq profiles around *ZIC2* gene in NT2-D1 and TIG-3 cells. The heat-map underneath ChIP-seq profiles shows the number of CpG nucleotides within 100bp sliding window (ranging from dark blue=0 to bright yellow=15). Here and in other figures, genes above the coordinate scale (in GRCh38/hg38 genomic release) are transcribed from left to right, genes below the coordinate scale are transcribed from right to left. The fragment used for transgenic experiment in (**E**) is indicated by red rectangle, with genomic position of ChIP-qPCR amplicon marked by vertical black line. **E.** ChIP-qPCR experiments indicate that the DNA underneath MEL18 peak downstream of the *ZIC2* gene generates new binding site for CBX2, MEL18, BMI1 and RING2 when integrated elsewhere in the genome. The transgenic DNA (*ZIC2* tran amplicon) immunoprecipitates as efficiently or better as corresponding DNA at native location (*ZIC2* end amplicon) or a positive control (*ALX4* locus, + control amplicon). Here and on all the following figures, the bar-plots show the average of three independent ChIP experiments performed with three independently prepared batches of chromatin. Circles show individual experimental results and whiskers show standard error of the mean. The CBX2, BMI1, MEL18 and RING2 ChIP signals for transgenic *ZIC2* amplicons are significantly higher (p < 0.05, unpaired, one-sided t-test) than that for the negative control (gene desert region). For all amplicons, the yields of the mock (no antibody) control ChIPs ranged from 0.008 to 0.07 % of input.

Like the *CCND2* PTE, the DNA underneath the *ZIC2* MEL18-bound peak is CpG-poor but surrounded by CpG-islands (Figure 1D). RT-qPCR analysis showed that, similar to *CCND2*, the *ZIC2* gene is highly transcribed in NT2-D1 cells but transcriptionally inactive in TIG-3 cells. (Figure 1C). Further resembling the *CCND2* case, in TIG-3 cells, the binding profiles of MEL18 and SUZ12 profiles differ. The MEL18 signal displays a major peak over the narrow region bound in NT2-D1 cells, with three smaller peaks further upstream. In contrast, the SUZ12 ChIP-seq signals are offset towards the CpG islands (Figures 1D, S1). Taken together, our observations fit the hypothesis that the DNA underneath the *ZIC2* MEL18-bound region contains a PTE. If so, we expect the corresponding DNA fragment to generate new PRC1 binding sites when integrated elsewhere in the genome. To test this, we cloned the corresponding 1.8kb DNA fragment into a lentiviral vector and integrated it back into the genome of NT2-D1 cells. To distinguish the transgenic and endogenous copies, we replaced a small stretch of nucleotides within the cloned DNA fragment to create an annealing site for the transgene-specific PCR primer (Figure S2, S3). ChIP-qPCR analysis indicates that the hypothetical PTE does generate new binding sites for MEL18, BMI1, CBX2 and RING2 (Figure 1E) when integrated elsewhere in the genome. Similar to the *CCND2* PTE, the transgenic *ZIC2* PTE is immunoprecipitatied with antibodies against SUZ12 just above the background level.

Taken together, our observations argue that screening ChIP-seq profiles for strong PRC1 signals coupled to low signals for PRC2 and high transcriptional activity of the nearby gene is a viable, although inefficient, strategy to find new PTEs. They also confirm our earlier observation that PRC1 and PRC2 may preferentially bind nucleotide sequences in different parts of the gene (Cameron et al., 2018).

### Discrete high-amplitude ChIP-seq peaks mark PRC1 tethering elements

Most of what we know about genomic binding of mammalian PRC1 and PRC2 comes from studies of mouse embryonic stem cells. There, the two complexes bind similarly and no sites stand out as potential PRC1 tethering elements (Fursova et al., 2019; Healy et al., 2019; Tamburri et al., 2020). Our ChIP-seq profiles from NT2-D1 and TIG-3 cell lines look different. At many genes repressed by Polycomb mechanisms (for example *ZIC2* in TIG-3 cells, Figure 1D), PRC1 ChIP-seq profiles have distinct sharp peaks. Could these ChIP-seq peaks mark high-occupancy PRC1 binding elements?

To evaluate this possibility, we searched the NT2-D1 MEL18 ChIP-seq profiles for discrete peaks. Briefly, we smoothed the ChIP-seq profiles and identified local maxima with amplitudes exceeding selected cutoffs (see Materials and Methods for detailed description of the peak-calling algorithm). Using this procedure, we identified 1014 peaks in the profile from the first MEL18 ChIP-seq experiment and 1152 peaks in the profile from the replicate experiment. We expect a genuine tethering element to be marked by a discrete MEL18 peak in each of the replicate profiles. In theory, corresponding peaks from replicate experiments should have the same genomic positions and ChIP-seq signal strength. In practice, both will differ, at least slightly, due to experimental variance. We therefore inspected peak sets from the two replicate MEL18 ChIP-seq experiments for pairs with the closest genomic positions. The survey of such pairs indicates that 84% of them have the peaks located less than 500bp apart, 61% - less than 250bp and in 33% of the cases, they reside as close as 100bp. As expected for peaks that represent the same MEL18 binding site, their ChIP-seq signal scores correlate (Figure S4B). Based on this, we assigned median genomic positions between paired peaks as centers of discrete MEL18 binding sites (potential PTEs) and used the distances between the corresponding paired peaks to reflect our confidence in each location (Figure 2A).

**Figure 2.**
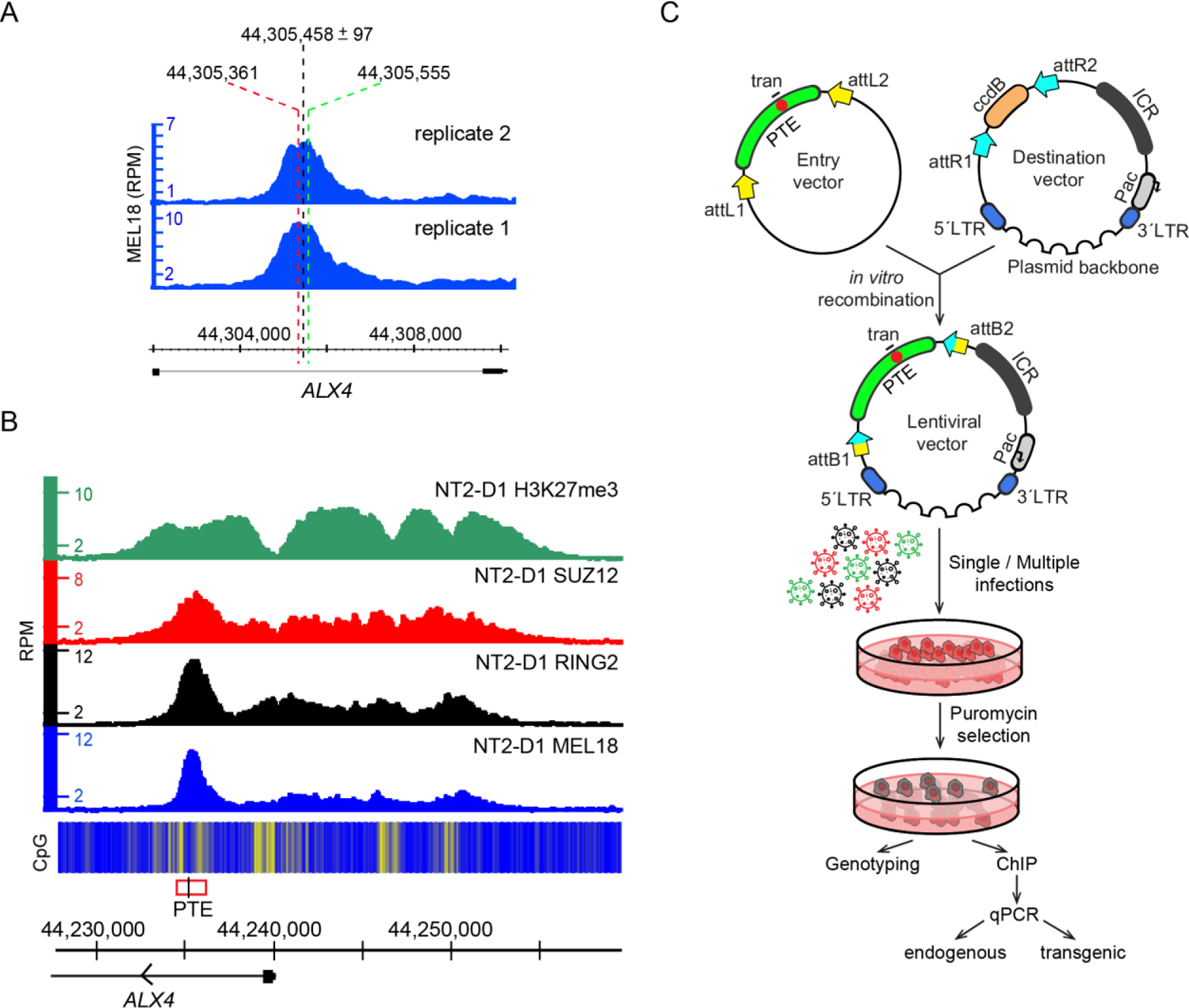
Distinct peaks mark PRC1 tethering elements. **A.** Assigning genomic positions of candidate PTEs. A screen-shot of replicate MEL18 ChIP-seq profiles around *ALX4* PTE from (**B**). Positions of corresponding MEL18 peak summits (marked with red and green dashed lines) deviate by 194bp due to experimental variance. The median genomic position between paired peaks is assigned as the center of candidate *ALX4* PTE. The half distance between the peaks estimates the accuracy of the candidate PTE location. **B.** H3K27me3, SUZ12, RING2 and MEL18 ChIP-seq profiles over *ALX4* locus, which harbors a candidate PTE (red rectangle, ChIP-qPCR amplicon indicated with black line) marked by obvious MEL18 peak. The heat-map underneath ChIP-seq profiles shows the number of CpG nucleotides within 100bp sliding window (ranging from dark blue=0 to bright yellow=15). **C.** The outline of the transgenic assay. To increase the throughput of the assay, the cloning procedure was modified to include Gateway attL/attR recombination (yellow and blue arrows). Cells were transduced with pools of up to six different lentiviral constructs. As in earlier experiments, a small stretch of nucleotides (red circle) within the cloned fragment (green arch) was replaced to distinguish the transgenic and endogenous copies.

We then split discrete MEL18 binding sites into quartiles based on the associated MEL18 ChIP-signal scores and selected six regions from the upper quartile (Q4). The selected regions had location accuracy of 168bp or better (obvious high and sharp peaks, Figures 2B, S5). After cloning the corresponding 1.4-2kb DNA fragments into a lentiviral vector, we integrated them back into the genome of NT2-D1 cells (Figure 2C, Figure S6). ChIP-qPCR indicates that all selected regions generate new PRC1 binding sites when integrated elsewhere in the genome of NT2-D1 cells (Figure 3A-B). The corresponding transgenic and endogenous regions display ChIP signals of similar strength. Taken together with results of transgenic analyses of *CCND2* (Cameron et al., 2018) and *ZIC2* elements, both of which correspond to well defined (± 35bp) upper quartile (Q4) MEL18 peaks, our observations argue that DNA fragments centered on sharp (± 300bp location accuracy) Q4 MEL18 peaks represent the high-occupancy PTEs.

**Figure 3.**
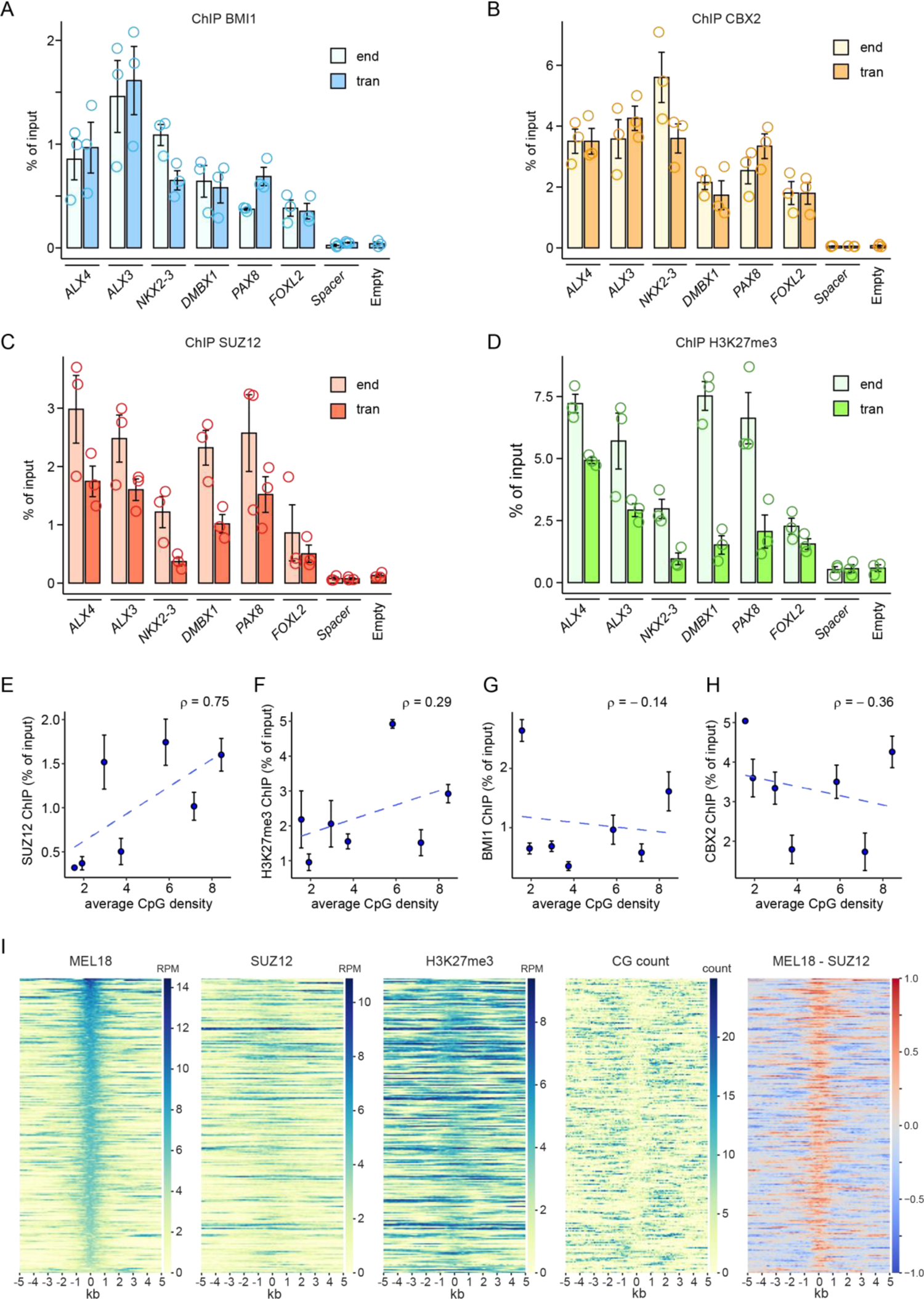
MEL18 Q4 peaks generate new PRC1-binding sites. ChIP-qPCR experiments indicate that DNAs of the PTEs from *ALX4*, *ALX3*, *NKX2.3*, *DMBX1, PAX8* and *FOXL2* loci generate new binding sites for BMI1 (**A**) and CBX2 (**B**) when integrated elsewhere in the genome. The BMI1 and CBX2 ChIP signals for all transgenic fragments (tran) are similar to those at endogenous locations (end). Both are significantly higher (p < 0.05, unpaired, one-sided t-test) than those for negative controls: the empty vector (Empty) and the gene desert region (Spacer). Note that the PCR primers to amplify the gene desert region do not discriminate between the transgenic and endogenous copies. Because the endogenous copy does not bind PRC1 or PRC2, any elevated ChIP signal in the transgenic cells (tran) would come from the transgene. For clarity, here and in Figures 4 and 6 immunoprecipitation yields for the endogenous *ALX4* amplicon (positive control) and endogenous Spacer amplicon are shown only for one of the transgenic cell lines. The values for other cell lines are comparable. Immunoprecipitation with the antibodies against SUZ12 (**C**) enriches for all transgenic PTEs (p < 0.05, unpaired, one-sided t-test). *ALX4*, *ALX3*, and *FOXL2* are also significantly enriched by ChIP with the antibodies against H3K27me3 (**D**). For all amplicons, the yields of the mock (no antibody) control ChIP reactions ranged from 0.0002 to 0.036 % of input. Scatter plots comparing the average CpG density to the average ChIP-qPCR signals for SUZ12 (**E**), H3K27me3 (**F**), BMI1 (**G**) and CBX2 (**H**) at transgenic PTEs from (**A**-**D**) and *ZIC2* (Figure 1E). Whiskers show the standard error of the mean. (**I**) Heat-map representations of ChIP-seq signals, CpG counts and MEL18-SUZ12 ChIP-seq signal differences (rightmost panel) around discrete MEL18 peaks indicate that PRC1 and PRC2 binding profiles differ.

### The binding of PRC1 and PRC2 is not strictly linked

Unlike PTEs from *CCND2* and *ZIC2* genes, some of the transgenic PTEs above are also strongly immunoprecipitated with antibodies against SUZ12 and H3K27me3 even when integrated elsewhere in the genome (Figure 3C-D). Consistent with previous observations (Hojfeldt et al., 2018; Lynch et al., 2012; Mendenhall et al., 2010), SUZ12 ChIP signal at transgenic PTEs positively correlates with their CpG content (Figure 3E). Weak positive correlation with CpG content is also seen for H3K27me3 signal (Figure 3F) but not for BMI1 and CBX2 (Figure 3G-H). These observations suggest that PRC1 and PRC2 binding is driven by distinct sequence features and that these features may be uncoupled or “intermixed”.

To explore the relation between PRC1 and PRC2 binding further, we compared ChIP-seq profiles for MEL18, SUZ12, H3K27me3 and CpG di-nucleotide distributions around all putative high-occupancy PTEs (Q4 MEL18 peaks). As illustrated by Figure 3I, in most instances, SUZ12 ChIP signals show no discrete peaks corresponding to those of MEL18. SUZ12 binding is broader, often shifted to the sides and correlated to higher CpG occurrence around PTEs.

Prompted by this observation, we asked whether the binding of PRC2 and associated H3K27me3 implies the co-binding of PRC1 at these sites or nearby. Comparison of genomic regions significantly enriched by ChIP with antibodies against SUZ12 and H3K27me3 to those enriched by immunoprecipitation with antibodies against MEL18 and RING2 indicates that this is not always the case. Thus, we detected 1628 regions bound by catalytically active PRC2 but lacking significant ChIP-seq signals for MEL18 within 10kb distance. At MEL18-free regions, the SUZ12 ChIP-seq signal is generally weaker compared to cases when SUZ12 binds next to discrete MEL18 peaks (Figure 4A). Nevertheless, some of them display SUZ12 ChIP-seq signal comparable to that next to high-occupancy PTEs. (Figure 4A, D). In both kinds of regions, the strength of the SUZ12 signal moderately (ρ ≈ 0.5) correlates with their CpG content. The correlation is weaker than that at the PTE transgenes suggesting that endogenous chromatin context influences PRC2 binding. Regardless of whether MEL18 is bound in the vicinity, the genes closest to SUZ12 bound regions tend to have low transcriptional output. (Figure 4C).

**Figure 4.**
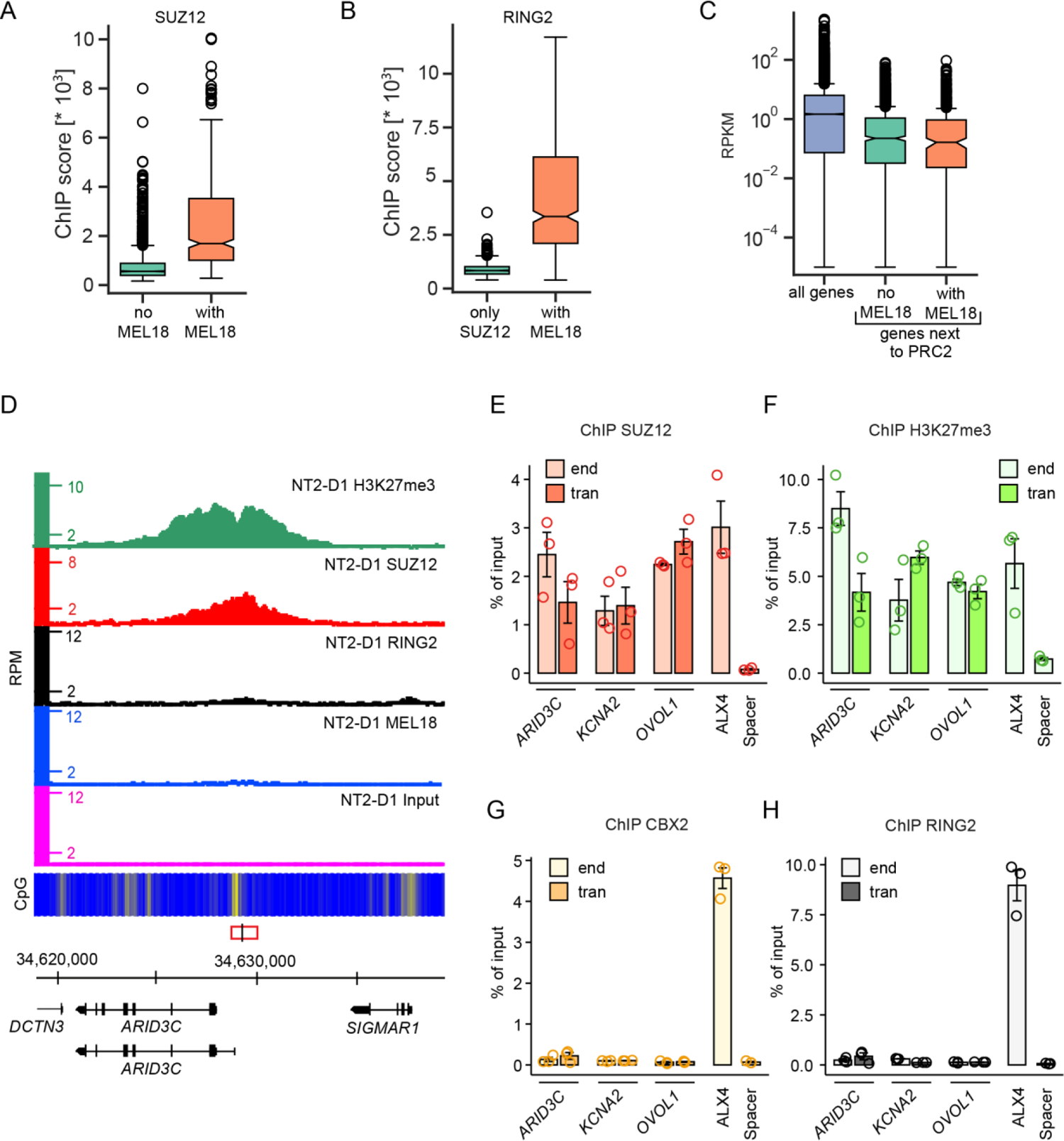
Uncoupled binding of PRC2 and PRC1. **A.** Box-plots of SUZ12 ChIP-seq signal scores at PRC1-free SUZ12 binding sites and sites with a discrete MEL18 peak nearby. **B.** Box-plots of RING2 ChIP-seq signal scores at PRC1-free SUZ12 binding sites and within discrete MEL18 peaks. **C.** Using data from (Li et al., 2019), transcriptional activity of TSSs throughout the genome is compared to that of the TSSs closest to MEL18-free SUZ12 binding sites or SUZ12 binding sites with a discrete MEL18 peak nearby. **D.** Screen-shot of the MEL18, RING2, SUZ12, H3K27me3 and Input ChIP-seq profiles around *ARID3C* gene in NT2-D1 cells. The fragment used for transgenic experiments is indicated by the red rectangle, with genomic position of ChIP-qPCR amplicon marked by vertical black line. Profiles show disproportionate signals for SUZ12 and H3K27me3 compared to MEL18 and RING2. ChIP-qPCR experiments indicate that the CpG-rich DNA fragments underneath SUZ12 peaks at *ARID3C*, *KCNA2* and *OVOL1* retain the ability to bind SUZ12 (**E**) and become tri-methylated at H3K27 (**F**) when integrated elsewhere in the genome. However, they do not generate new binding sites for CBX2 (**G**) and RING2 (**H**). For all amplicons, the yields of the mock (no antibody) control ChIPs ranged from 0.0001 to 0.05 % of input.

Is the information included in the nucleotide sequences of the MEL18-free PRC2-bound regions sufficient to generate new PRC2 binding sites? To address this question, we generated three lentiviral transgenic constructs containing such regions from *ARID3C*, *KCNA2* and *OVOL1* loci (Figures 4D, S7A-B) and integrated them elsewhere in the genome of the NT2-D1 cells. Strikingly, all three constructs are immunoprecipitated with antibodies against SUZ12 and H3K27me3 but not with antibodies against CBX2, BMI1 or RING2 (Figure 4E-H, S7C-F, S8). This suggests that MEL18-free PRC2-bound regions are sufficient to tether PRC2.

Although not confidently identified by qPCR at the three selected sites, ChIP-seq experiments detect RING2 at some of the MEL18-free PRC2-bound regions. The RING2 ChIP-seq signal at these regions is much weaker than that at discrete MEL18 binding peaks (Figure 4B). We speculate that these weak signals represent the binding of RING-RYBP complexes. Alternatively, they may represent exceedingly weak or transient binding of canonical PRC1 detectable only by ChIP with antibodies against RING2. Regardless, our observations argue that, in the NT2-D1 cells, PRC2 binding and associated tri-methylation of H3K27 is not sufficient to tether canonical PRC1 and, inversely, that the catalytically active PRC2 can be tethered to a locus without appreciable co-binding of PRC1. Overall, it appears that PRC1 and PRC2 binding is driven by distinct DNA sequence determinants. These determinants may be uncoupled or intermixed to a variable degree within a single DNA stretch.

### High-occupancy PTEs repress transcription of the nearby genes

Distinct high-amplitude PRC1 ChIP-seq peaks mark high-occupancy PTEs sufficient to generate new PRC1 binding sites when integrated elsewhere in the genome of the NT2-D1 cells. Are these elements necessary for PRC1 binding at their endogenous locations and, more importantly, do they repress transcription? To answer these questions, we sought to delete PTEs from genes bound by both PRC1 and PRC2 using CRISPR/Cas9 – mediated genome editing. Our initial attempts to delete PTEs in NT2-D1 and TIG-3 cells where marred by technical difficulties. NT2-D1 cells are hypotriploid (Andrews, 1988) and TIG-3 cells are extremely difficult to clone. To facilitate the genome editing, we turned to the nearly haploid human myeloid leukaemia cell line KBM7-1-55-S2-24 (clonal derivative F10) (Kotecki et al., 1999), thereafter referred to as F10. These cells have only a single copy of a chromosomal site to edit and are easy to clone. First, we mapped the distribution of BMI1, RING2, SUZ12 and H3K27me3 throughout the genome of these cells using ChIP-seq. It showed that, in F10 cells, *CCND2*, *ZIC2* and *ALX3* genes are bound by PRC1, PRC2 and H3K27me3 (Figure 5A-C) and likely repressed by the Polycomb system. We therefore transfected F10 cells with mixtures of the Cas9 protein and gRNAs targeting *CCND2*, *ZIC2* or *ALX3* PTEs respectively and derived four clonal cell lines with precise deletion of the corresponding PTE from each treated cell pool (Figure S9).

**Figure 5.**
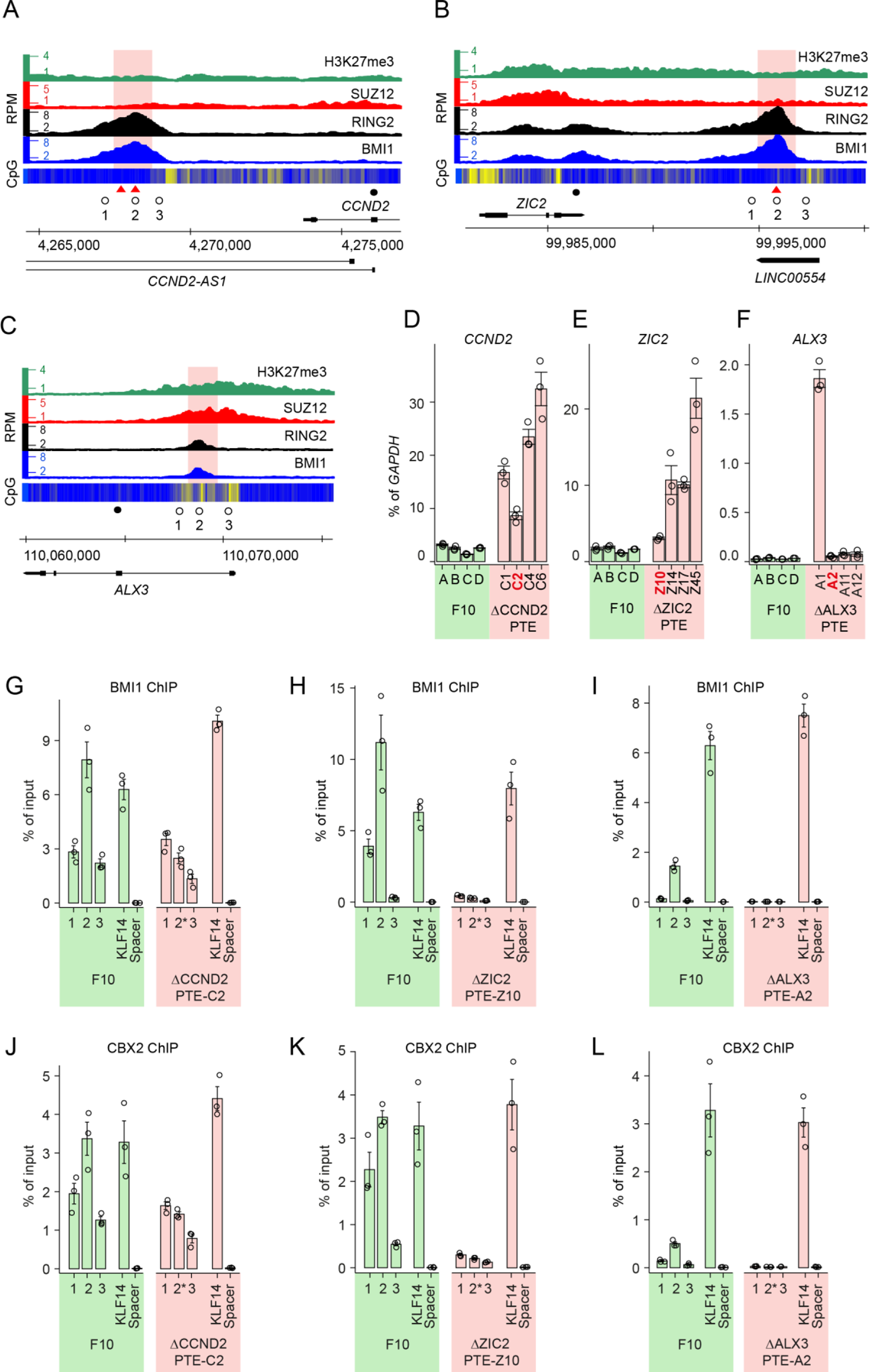
High-occupancy PTEs repress transcription. Screen-shots of BMI1, RING2 SUZ12 and H3K27me3 profiles throughout *CCND2* (**A**), *ZIC2* (**B**) and *ALX3* (**C**) loci in F10 cells. Open circles mark positions of the PCR amplicons used for ChIP-qPCR, filled circles mark positions of the RT-qPCR amplicons. Red triangles indicate positions of the AAACGAAA motifs. The pink shading marks DNA stretches deleted by CRISPR/Cas9-mediated editing. RT-qPCR measurements of *CCND2* (**D**), *ZIC2* (**E**) and *ALX3* (**F**) transcription in four clones of unedited F10 cells (green bars) and four clones of F10 cells with corresponding deletions (pink bars). The clones used for ChIP are marked in red. ChIP-qPCR with antibodies against BMI1 (**G**-**I**) and CBX2 (**J**-**L**). Immunoprecipitation of *KLF14* and an intergenic region (Spacer) were used as positive and negative controls, respectively. Note that 2* amplicons span the deletion breakpoints.

When comparing parental cell line to its genetically manipulated clonal derivative a difference in transcriptional output of a gene may stem from genetic manipulation (i.e. PTE deletion) or from individual cell-to-cell variation already present in the population of parental cells and fixed by cloning. To control for the latter, we derived four cell lines by expansion of single cells from the original F10 population (lines F10A, F10B, F10C, F10D). The RT-qPCR measurements show little variation in transcript levels of *CCND2*, *ZIC2* and *ALX3* between the control cell lines (Figure 5D-F). In contrast, all four clonal derivatives lacking the *CCND2* or *ZIC2* PTEs display variable but significant increase in the RNA levels of the corresponding genes (Figure 5D-E). Only one line with deleted *ALX3* PTE shows higher *ALX3* transcript abundance (Figure 5F). We speculate that F10 cells express the subthreshold quantities of transcription factors required to activate *CCND2* and *ZIC2*. These quantities are not sufficient to drive transcription effectively in the presence of PTEs but become so when PTEs are deleted. In the case for *ALX3,* whose transcription in the unedited F10 cells is essentially undetectable, the levels of corresponding transcriptional activators must be very low. Taken together, our observations argue that deletion of high-occupancy PTEs leads to stochastic increase in transcription of the closest genes and, therefore, the high-occupancy PTEs are repressive elements.

When PTEs are removed, the high ChIP-signals for BMI1 and CBX2 across corresponding deletion breakpoints disappear (Figures 5G-L). This indicates that the *CCND2*, *ZIC2* and *ALX3* PTEs are required for efficient tethering of PRC1 at these loci. Interestingly, deletion of PTEs from *ALX3* and *ZIC2* loci causes nearly complete loss of PRC1 ChIP signals including those in the flanking regions (Figures 5H, I, K, L). This suggests that, in some loci, PTEs are the master determinants of PRC1 binding. In contrast, at *CCND2*, the weaker PRC1 ChIP signals within the flanking regions are unaffected by the PTE deletion (Figures 5G, J). It appears that, in addition to the PTE required for high occupancy binding, *CCND2* contains additional DNA features capable to maintain some degree of PRC1 tethering.

### PRC1 tethering is not limited to high-occupancy PTEs

DNA sequences around *CCND2* PTEs provide some degree of PRC1 tethering. Therefore, we wondered whether weaker (Q3-Q2) MEL18 ChIP-seq peaks from our data set could do the same. More specifically, we wished to ask whether DNA fragments underneath such MEL18 peaks could generate new PRC1 binding sites when integrated elsewhere in the genome.

To address this question we selected eight discrete (position accuracy better than ±167bp) MEL18 peaks from Q3-Q2 group (Figures S10, S11) and re-introduced them in the genome of the NT2-D1 cells (Figure S12). ChIP analysis of the resulted transgenic cell lines revealed two trends. First, all transgenic fragments displayed significantly stronger precipitation with anti-BMI1 and anti-CBX antibodies (p-value < 0.05, unpaired, one-sided t-test) compared to the negative control (Figure 6A-B). ChIPs with antibodies against SUZ12 and H3K27me3 were more variable, and some amplicons were not significantly enriched compared to the negative control (Figure 6C-D). Second, in contrast to the high-occupancy PTEs (Figures 3A-B, 6E-F), the signals for the Q3-Q2 fragments at endogenous location and within the transgene were not positively correlated (Figures 6A-B, 6G-H).

**Figure 6.**
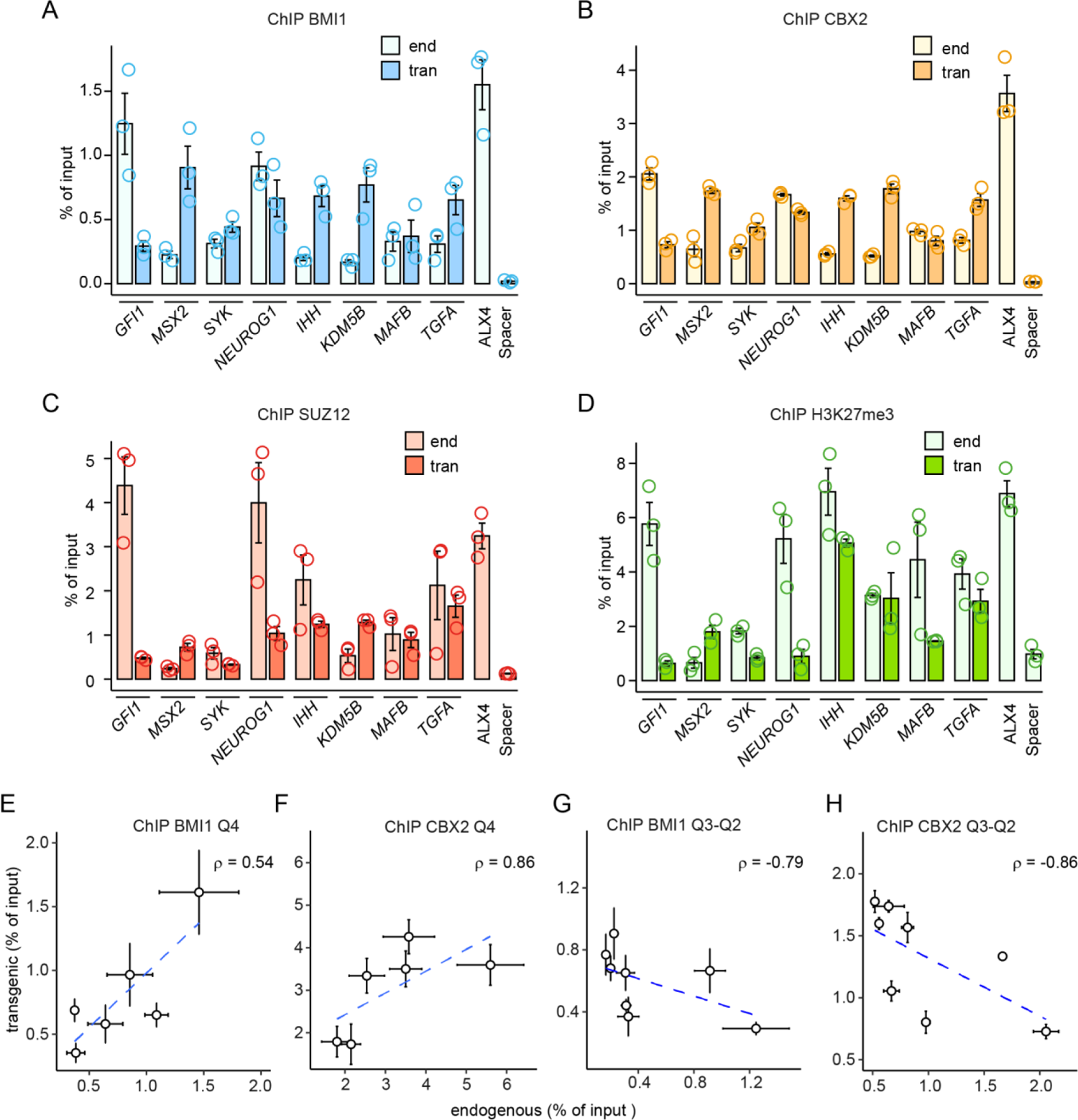
PRC1 tethering by low-occupancy sites. ChIP-qPCR experiments indicate that DNA fragments underneath selected Q3 and Q2 MEL18 peaks always generate new binding sites for BMI1 (**A**) and CBX2 (**B**) when integrated elsewhere in the genome. Sometimes they also generate new binding sites for SUZ12 (**C**) and H3K27me3 (**D**). Immunoprecipitation of *ALX4* and an intergenic region (Spacer) were used as positive and negative controls, respectively. Comparison of BMI1 (**E**, **G**) and CBX2 (**F**, **H**) ChIP-qPCR signals at transgenic and endogenous locations indicate that those, corresponding to the high-occupancy PTEs, correlate but those, corresponding to the low occupancy MEL18 peaks, do not. The data for high-occupancy PTEs is from Figure 3A-B. The whiskers show the standard error of the mean for three independent experiments.

We take it to indicate that the majority of the DNA fragments underneath Q2 and Q3 MEL18 peaks can tether PRC1 when integrated elsewhere in the genome. However, compared to that of the high occupancy PTEs, the tethering ability is more dependent on the chromatin context. The information contained within the DNA underneath the Q2 and Q3 MEL18 peaks is essential for PRC1 tethering as neither the “empty” backbone nor the constructs containing MEL18-free PRC2-bound CpG-islands or Spacer fragments generate PRC1 binding sites. Yet, our lentiviral transgenic system appears to favor tethering as some of the Q3-Q2 PTEs display stronger PRC1 ChIP-signals inside the transgenes compared to their endogenous locations.

### DNA sequence determinants of PRC1 tethering

The information contained within PTEs is in some way encoded in their DNA sequence. What sequence features define this code? To address this issue we first examined the overall nucleotide composition of the DNA underneath and around discrete MEL18 peaks. As illustrated by Figure 7A, this DNA is enriched in AA and TT di-nucleotides and is flanked by the CpG-rich nucleotide sequences. These nucleotide sequence features are most prominent within MEL18 peaks that display the highest ChIP-signal scores and become progressively less distinct in peaks with lower MEL18 ChIP-signals (Figure S13).

**Figure 7.**
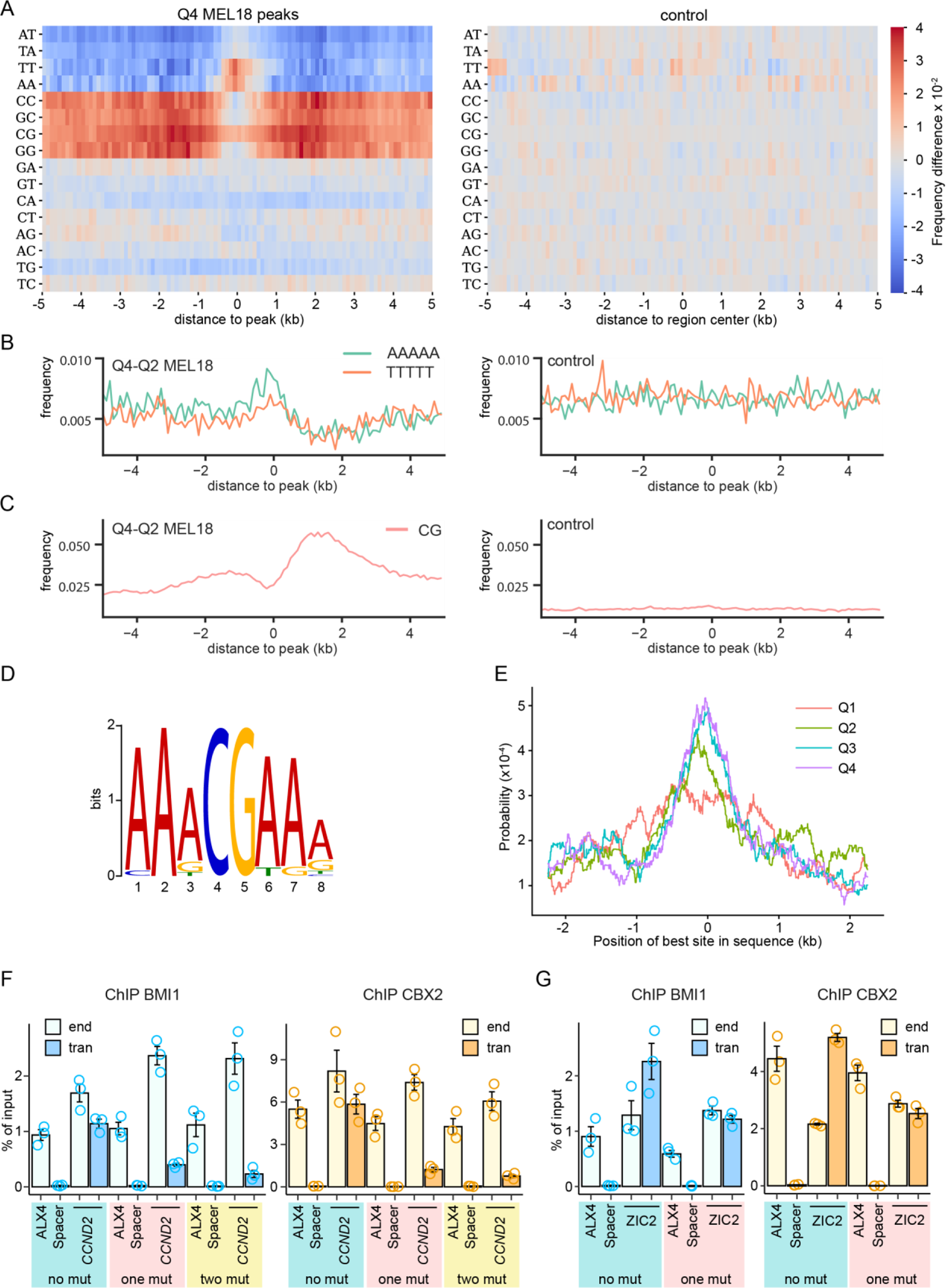
DNA sequence features of PTEs. **A.** Heat-map representation of the di-nucleotide frequency differences around high-occupancy (Q4) MEL18 peaks and randomly selected control sequences that do not bind PRC1 or PRC2. The control map is produced by calculating the frequency difference between randomly divided sequences of the control group. **B.** The frequency of poly(dA)_5_ and poly(dT)_5_ within 100bp windows around MEL18 peaks and within control DNA sequences. **C.** CpG frequency for the same DNA sequences as in (**B)**. **D.** Web logo representation of the “AAACGAAA” motif. **E.** Probabilities to find the best match to the “AAACGAAA” motif around MEL18 peaks of different occupancy (ChIP-seq score quartiles). The results of ChIP-qPCR with chromatin from NT2-D1 cells transduced with various *CCND2* PTE (**F**) and *ZIC2* PTE transgenes (**G**).

Although the DNA underneath the MEL18 peaks is enriched in AA and TT di-nucleotides, its AT or TA content remains close to the genomic average (Figures 7A, S13). This suggests that the presence of poly(dA:dT) nucleotides, rather than general AT-richness, in some way helps to tether PRC1. Certain budding yeast promoters have the strand-biased poly(dA) distribution (Wu and Li, 2010), apparently, to take advantage of the directional nucleosome sliding activity of the RSC chromatin remodeling complexes, which is stimulated by the homopolymeric poly(dA:dT) tracts (Krietenstein et al., 2016; Lorch et al., 2014). To explore this resemblance, we oriented the nucleotide sequences underneath the high-occupancy (Q4) MEL18 peaks such that the TSS of the closest gene was always to the right of the peak. Plotting the frequency of poly(dA)_5_ and poly(dT)_5_ tracts across the oriented DNA sequences revealed two noticeable biases (Figure 7B). First, the DNA underneath MEL18 peaks is enriched in poly(dA)_5_ tracts. Second, the distributions of poly(dA)_5_ and poly(dT)_5_ tracts around the MEL18 peaks differ. Both kinds of tracts become less frequent, compared to the genomic average, in the DNA to the right of the peaks, presumably reflecting its elevated CG-content (Figure 7C). In contrast, while the frequency of the poly(dT)_5_ tracts within DNA to the left of the MEL18 peaks follows the same trend, the frequency of the poly(dA)_5_ tracts remains high. It appears that selection pressure prevents the poly(dA)_5_ but not the poly(dT)_5_ tracts from being depleted within this CG-rich DNA. The biases are also evident for the DNA under the high-occupancy MEL18/BMI1 peaks identified by ChIP-seq with TIG-3 and F10 cells. The RSC complex (known as PBAF in mammals) is evolutionarily conserved (Clapier and Cairns, 2009). Conceivably, PRC1 tethering by PTEs benefits from the directional sliding of nucleosomes.

Nucleosome displacement may expose binding sites recognized by the sequence-specific DNA binding proteins that act as adaptors to tether of PRC1. To investigate this possibility, we combined the STREAM (Bailey, 2021) and CentriMo (Bailey and Machanick, 2012) algorithms implemented in MEME software suite (Bailey et al., 2015) to identify sequence motifs enriched in the DNA underneath MEL18 peaks and biased towards their summits. Analysis of the 5kb DNA fragments centered on high-occupancy (Q4) MEL18 peaks revealed one such motif that we dubbed “AAACGAAA” (Figure 7D). The motif also showed significant preference towards the summits of Q3 and Q2 MEL18 peaks. (Figure 7E).

Comparison with experimentally defined transcription factor binding sites from JASPAR (Castro-Mondragon et al., 2022) and HOCOMOCO (Kulakovskiy et al., 2018) databases revealed no candidate proteins to bind the “AAACGAAA” motif. Its sequence resembles previously reported “CGA” motif, which we found enriched in the MEL18/BMI1 binding sites on human chromosomes 8, 11, and 12 (Cameron et al., 2018). We posit that the “AAACGAAA” motif is a better-defined version of “CGA” due to a more numerous and accurately mapped PRC1 binding sites employed in the present analysis.

The *CCND2* PTE contains one AAACGAAA nucleotide sequence, the highest-scoring match to the motif’s position weight matrix, located directly underneath of the MEL18 peak summit (Figure 5A). In addition, it harbours the second best match, AATCGAAA, just at the edge of the 1kb fragment that generates new PRC1 binding site when integrated elsewhere. Converting the best matching AAACGAAA sequence to AAAAAAAA leads to the four-fold reduced immunoprecipitation of the transgenic *CCND2* PTE with antibodies against BMI1 and CBX2 (Figure 7F, S14). Simultaneous conversion of the second best-matching motif has no additional impact on the PRC1 tethering (Figure 7F). The *ZIC2* PTE has no best-matching instances of the motif but has the AATCGAAA sequence located just underneath the MEL18 peak summit (Figure 5B). Conversion of this sequence to AAAAAAAA reduces the immunoprecipitation of the transgenic PTE two-fold (Figure 7G, S14). Taken together, our observations argue that the “AAACGAAA” motif plays significant role in tethering PRC1 to PTEs. Additional studies will be needed to uncover the molecule that recognizes it.

### PTEs tether PRC1 to developmental genes

In the NT2-D1 cells, our algorithm identifies 913 MEL18 peaks of which 396 we consider as putative high-confidence PTEs because they have high (Q4 or Q3) MEL18 ChIP-seq signal score and the position defined with accuracy better than ± 301bp (Table S1). What kind of genes are regulated by these PTEs? To address this question we searched for the nearest Transcription Start Sites (TSS) to the left and to the right of each PTE and asked if these TSS are significantly enriched by immunoprecipitation with antibodies against H3K27me3. In 75% of cases, the chromatin of the closer of the two TSS was enriched in H3K27me3. We took this to indicate that the corresponding gene is the most likely target of the PTE. This is a conservative approach. At some loci, a PTE may affect several genes, which we did not consider for further analyses. In few cases (4%), the closer of the two TSS was not enriched in H3K27me3, but the other one was. In those instances, the gene corresponding to the latter was designated as the likely PTE target. In 21% of the cases, which included *CCND2* and *ZIC2* PTEs, neither of the nearby TSSs was enriched by H3K27me3 ChIP. In those cases, the gene with the TSS closest to the PTE was designated as the likely target.

The putative high-confidence PTEs are shared between 277 genes. Conversely, 29% of these genes are associated with more than one (from 2 to 5) of them. The majority of the genes have the closest PTE at a distance from 0.5kb to 7kb (Figure S15). Analysis of the gene ontology terms associated with the PTE-equipped genes indicate that many of them are regulators of cell fate commitment, pattern specification, embryonic organ development, neuron fate commitment and forebrain development (Figure 8A). Many of these genes encode for transcription factors and 39% are linked to heritable human syndromes (Table S2). Using the same algorithm, we identified 125 genes regulated by putative high-confidence PTEs in TIG-3 cells. Our ChIP-seq analyses identified fewer regions enriched by MEL18 ChIP in these cells, which explains the smaller number of putative high-confidence PTEs and target genes. 62.4% of these genes are common between the two cell lines. Despite only partial overlap, the PTE associated genes from TIG-3 cells are also enriched for regulators of cell fate commitment, pattern specification and embryonic organ development (Figure S16). In addition, they include regulators of skeletal and muscle tissue development, not overrepresented in the NT2-D1 gene set, which may reflect the fibroblast origin of TIG-3 cells (Bracken et al., 2006). When compared to published RNA-seq data (Li et al., 2019), 82% of the PTE-equipped genes from NT2-D1 cells have no or little transcriptional activity (ranked below median transcription level, hereafter referred to as genes of transcription groups G1-G2). 10% of the genes display moderate transcription (genes with RNA-seq signal within the third quartile, transcription group G3) and only 7%, exemplified by the *CCND2* and *ZIC2* genes, are highly transcriptionally active (genes with RNA-seq signal from the upper quartile, transcription group G4). As expected, most of these genes have chromatin around TSS decorated with H3K27me3. The extent of H3K27 methylation inversely correlates with transcriptional activity and it appears to be progressively lost with increased transcription starting from regions immediately around TSSs (Figure 8B). The latter may reflect the increased nucleosome exchange that accompanies transcription initiation or the necessity to remove H3K27me3 at the TSS in order to commence transcription.

**Figure 8.**
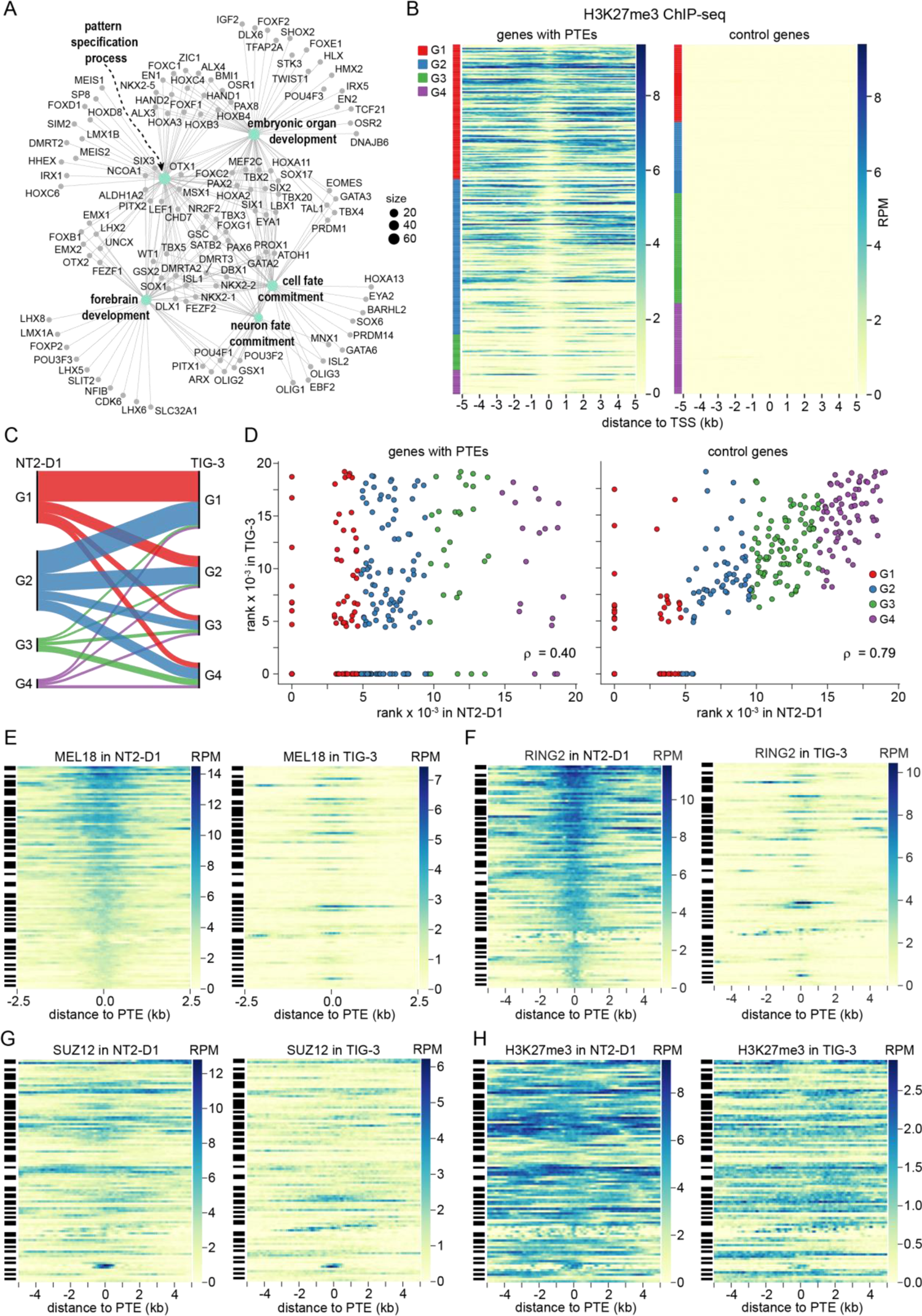
PTE regulated genes. **A.** Genes regulated by the PTEs in the NT2-D1 cells grouped in a network by overrepresented GO terms. **B.** ChIP-seq signals for H3K27me3 around TSSs of PTE-equipped and control genes ranked by their transcriptional activity (the least transcribed at the top, the most transcribed at the bottom). The color code at the left indicates corresponding transcriptional quartiles. **C.** Sankey diagram of transcriptional changes of the putative PTE-regulated genes between NT2-D1 and TIG-3 cells. **D.** Scatter plots of transcriptional activity ranks for putative PTE-regulated genes and control genes that do not bind any Polycomb complexes in NT2-D1 and TIG-3 cells. Heat-map representations of MEL18 (**E**), RING2 (**F**), SUZ12 (**G**) and H3K27me3 (**H**) ChIP-seq signal around PTEs of genes that remain transcriptionally inactive (belong to G1) in both NT2-D1 and TIG-3 cells. Streak-codes to the left of heat maps indicate PTEs belonging to the same locus. Note loci bound by Polycomb complexes and H3K27me3 in NT2-D1 and devoid of either in TIG-3 cells.

Judging from published RNA-seq data (Li et al., 2019; Yano et al., 2021), PTE-equipped genes with low transcriptional activity in NT2-D1 tend to stay in that state also in TIG-3 cells. However, some genes change to moderate or highly transcribed (Figure 8C). Conversely, some of the genes highly transcribed in NT2-D1 cells (e.g. *CCND2* and *ZIC2*) become transcriptionally inactive in TIG-3 (Figure 8C). We note that the levels of transcriptional activity in NT2-D1 and TIG-3 cells for genes not regulated by Polycomb system are well correlated (ρ = 0.75; Figure 8D). In contrast, the transcriptional output from PTE-equipped genes is much less consistent (ρ = 0.4; Figure 8D) suggesting that these genes are predisposed to change their transcription more discretely. Epigenetic regulation by Polycomb system likely contributes to this property.

Transcriptional activity is known to antagonize PRC2 binding and H3K27 methylation by means that are not fully understood (Bowman et al., 2014; Holoch et al., 2021; Schwartz et al., 2010) and even PRC1 is sometimes lost from transcriptionally active *Drosophila* genes (Schwartz et al., 2006). However, factors other than those linked to transcription modulate the binding of PRC1 in human cells. Side-by-side comparison of PRC1 binding to genes that are transcribed at very low levels (belong to group G1) in both NT2-D1 and TIG-3 cells indicates that, in the latter, many PTEs are no longer immunoprecipitated with either anti-MEL18 or anti-RING2 antibodies (Figures 8E-F). In some instances, MEL18 and RING2 signals are lost from one PTE but remain above genomic average at other PTEs of the same gene. Yet, in 42% of the cases, all PTEs of a gene, which were occupied by MEL18 in NT2-D1 cells, display no immunoprecipitation with antibodies against MEL18 in TIG-3 cells. In one third of these genes, all PTEs in TIG-3 cells also lack detectable RING2 ChIP-seq signal. The ChIP-seq signals for MEL18 and RING2 in TIG-3 cells are generally lower compared to those in NT2-D1 cells. This is consistent with lower transcription of *MEL18* and *RING2* genes (Cameron et al., 2018) and may indicate generally lower abundance of PRC1 in TIG-3 cells. However, this does not fully account for the PRC1 loss. Thus, some of the PTEs that display very high MEL18 and RING2 ChIP-seq scores in NT2-D1 cells are no longer immunoprecipitated in TIG-3 cells, while some of those with weaker scores remain occupied in both cell lines. Consistent with the gene sets being transcriptionally inactive in both cell lines, most of the genes retain significant presence of SUZ12 and H3K27me3 in TIG-3 cells (Figures 8G-H).

Overall, our observations suggest that canonical PRC1 complexes do not bind PTEs of transcriptionally inactive target genes by default and additional processes “license” PTEs for binding. The extensive catalogue of human PTEs presented here provides a major new resource to discover these processes.

## DISCUSSION

Three central conclusions follow from the study presented here. First, many human developmental genes contain DNA elements necessary and sufficient for tethering canonical PRC1. Second, the binding of PRC1 and PRC2 to a regulated locus is not strictly linked and the presence of PRC2-catalyzed H3K27me3 is not enough for efficient PRC1 tethering. Third, the DNA features associated with PRC1 tethering differ from those that favour the retention of PRC2. Throughout the genome, the two kinds of sequence features mix in different proportions to yield a gamut of DNA elements that range from those tethering predominantly PRC1 to ones that can tether both PRC1 and PRC2.

The discovery of hundreds of PRC1 Tethering Elements puts to rest the question of whether DNA elements play a role in directing canonical PRC1 to human genes (Bauer et al., 2016; Owen and Davidovich, 2022). The emerging picture is similar to the paradigmatic targeting of Polycomb complexes by Polycomb Response Elements (PREs) of *Drosophila* but with instructive differences. Both organisms appear capable of tethering PRC2 to specific sites independently of PRC1 (Kahn et al., 2016). Yet, in *Drosophila*, the high-occupancy sites for the two complexes always coincide at PREs while, in human cells, PRC1 and PRC2 often prefer to bind different parts of the gene.

The distinction may stem from dissimilar primary structures of human and *Drosophila* genomes. Shaped by abundant CpG DNA methylation, the human genome is generally CpG-poor save for complex regulatory regions of tissue and cell-type specific genes (Deaton and Bird, 2011; Schubeler, 2015). These genes appear “automatically” marked for preferential binding by PRC2, which has a propensity to bind CpG-rich DNA (Hojfeldt et al., 2018; Li et al., 2017). When equipped with one or more PRC1 tethering elements (PTEs), such a gene is set for regulation by the Polycomb system. *Drosophila melanogaster*has lost the CpG methylation (Engelhardt et al., 2022) and, with it, the ability to distinguish the tissue and cell-type specific genes from the rest of the genome by their CpG-content. We speculate that flies had to evolve a tighter coordination in tethering of PRC2 and PRC1 (Kahn et al., 2016; Kang et al., 2015) to compensate for this loss.

Most of what we know about genomic binding of mammalian PRC1 and PRC2 is derived from studies of mouse embryonic stem cells. There PRC1 and PRC2 bind repressed genes broadly with no sites standing out as being highly occupied by PRC1 (Fursova et al., 2019; Healy et al., 2019; Tamburri et al., 2020). The findings presented here paint a different picture. While we cannot exclude that distinct ChIP-seq profiles of human PRC is a feature of this species, we think it is unlikely. For example, ChIP-qPCR mapping of PRC1 and PRC2 across the murine *Ccnd2* gene in NIH-3T3 mouse embryonic fibroblasts revealed distinct PRC1 peak orthologous to human *CCND2* PTE (Cameron et al., 2018). Moreover, the DNA underneath this peak generates new PRC1 binding site when introduced into the human genome (Cameron et al., 2018).

Some of the apparent differences may stem from the adjustment of ChIP-seq procedure where we omitted the size-selection step and, instead, fragmented immunoprecipitated DNA enzymatically prior to ligation of adapters for Illumina sequencing. In our hands, this makes immunoprecipitation profiles obtained by the next-generation sequencing more comparable to those derived from the same ChIP reaction by qPCR. Yet, we suspect that the major source of the discrepancy is inherent to mouse embryonic stem cells and comes from their unusually high levels of PRC2 (Kloet et al., 2016; Stafford et al., 2018) and PRC1. Systematic comparison of PRC1 and PRC2 abundance to their binding patterns will help to clarify this issue. Regardless, our observations suggest that the behaviour of the Polycomb system in mouse embryonic stem cells is not representative of all mammalian cell types.

Most contemporary models of mammalian Polycomb system assume that canonical PRC1 binds to repressed genes via interaction of its CBX subunit with H3K27me3. This assumption does not easily fit the observations presented here. In the NT2-D1, TIG-3 and F10 cells, the ChIP-seq profiles of canonical PRC1 and PRC2 differ and the presence of PRC2-catalyzed H3K27me3 is not sufficient to achieve PRC1 occupancy comparable to that at PTEs. It is worth to note that, in *Drosophila*, for which the H3K27me3 - CBX hierarchy was first proposed (Cao et al., 2002), PRC1 remains bound to PREs in cells depleted of PRC2 and H3K27 methylation (Kahn et al., 2016). To summarize, even though H3K27 methylation is essential for repression (Pengelly et al., 2013; Sankar et al., 2022), its mechanistic contribution remains to be fully understood.

The overrepresentation of oriented poly(dA) tracts within PTEs suggests that PRC1 tethering is somehow linked to chromatin remodeling by SWI/SNF (i.e. BAF or PBAF) complexes. In line with this notion, Weber and co-authors have found that rapid degradation of BRG1, the core ATPase subunit of BAF and PBAF complexes, leads to substantial loss of PRC1 from the most highly occupied binding sites in mouse embryonic stem cells (Weber et al., 2021). It would be interesting to investigate whether the same holds true for human cells and, if so, which of the two complexes, BAF or PBAF, is implicated. The necessity of chromatin remodeling would explain why canonical PRC1 complexes do not always bind PTEs of transcriptionally inactive genes.

The work presented here advances our understanding of how human PRC1 binds to specific genes. It also raises a number of new questions. Do PTEs vary in their ability to tether PRC1 complexes composed of different CBX or PHC paralogues? How do deletions of individual PTEs affect the expression of their cognate genes in the context of the whole organism? What fraction of PTE-equipped genes linked to heritable human syndromes has alleles that carry PTE deletions or duplications? Our catalogue of human PTEs provides a valuable new resource to address these questions. Equally significant, it presents an opportunity to design transgenic assays to disentangle individual contributions of PRC1 and PRC2 to the repression by the Polycomb system and to screen for factors that enable the PRC1 tethering.

## MATERIALS AND METHODS

### Cell culture

NTERA-2 (NT2-D1 ATCC® CRL-1973™), TIG-3 (see (Cameron et al., 2018) for more details), 293T (ATCC® CRL-3216™) and Hela (ATCC® CCL-2™) were cultured in high-glucose DMEM (Gibco) supplemented with 10% of heat inactivated foetal bovine serum (Sigma), penicillin/streptomycin (Gibco), and 1.5 g/liter sodium bicarbonate (Sigma) at 37 °C in an atmosphere of 5% CO2. KBM7-1-55-S2-24 cell line (Kotecki et al., 1999) was a gift from Dr. Brent Cochran (Tufts University). Thwese cells were cultured in Iscove’s Modified Dulbecco’s Medium (1X) (Gibco) supplemented with 15% of heat inactivated foetal bovine serum (Sigma), 1x penicillin/streptomycin (Gibco), and 3.024 g/liter sodium bicarbonate at 37 °C in an atmosphere of 5% CO2. Several sub clones were isolated from the original cell line by limited dilution and their karyotypes ecvaluated as follows. Cells were incubated with colchicine (0.1µg/ml) for 75 minutes, harvested, incubated in hypotonic solution (KCl 75mM) for 20 minutes at 37^0^C and fixed in methanol/glacial acetic acid (3:1). Metaphase spreads were analysed manually. Two sub clones (F10 and D4) with most similar karyotypes [25,XY,+8,t(9;22),add(19q)] were selected and the F10 sub clone was used for most experiments.

### Lentiviral transgenesis

Two transgenic strategies were used to test whether selected DNA fragments can autonomously recruit PRC1 and/or PRC2 when integrated elsewhere in the genome. Both employed the same approach to discriminate between the endogenous and the transgenic DNA copies. For all tested fragments, we identified small stretches of nucleotide sequences that were well enriched by ChIPs with antibodies against MEL18/BMI1 (for PTEs) or SUZ12 (for MEL18-free PRC2-bound regions) but showed little conservation within mammalian species. We then substituted four of those nucleotides in the transgenic copies to create an annealing site for transgene-specific PCR primers (see Figure S2).

The first transgenic strategy, used to test the *ZIC2* PTE (experiments shown on Figure 1D-E), is essentially the same as described by (Cameron et al., 2018). Briefly, the pLenti-ZIC2 PTE 1.8-kb construct was generated by digesting the pLenti-ICR-Puro vector (Cameron et al., 2018) with AfeI and PmlI and recombining it with two overlapping DNA fragments encompassing 1.8kb of the putative *ZIC2* PTE and with a four base pairs mutation for specific PCR primer. These two PCR fragments were amplified using ZIC2_1.8_PTE_F and ZIC2_trans_R primers, and ZIC2_trans_F and ZIC2_1.8_PTE_R primers, and human genomic DNA as a template. pLenti-ZIC2_mutTCG transgenic construct was generated the same way except that corresponding 1.8kb DNA fragment was PCR amplified using ZIC2_1.8_PTE_F and ZIC2_mutTCG_2.2 primers, and ZIC2_mutTCG_1.2 and ZIC2_1.8_PTE_R primers and using DNA of the pLenti-ZIC2 PTE 1.8-kb construct as a template. Similarly, for the generation of the pLenti_CCND2_PTE_2xCGmut two PCR fragments were amplified using CCND2_mutTCG_1.1 and CCND2_mutTCG_1.2 primers, and CCND2_mutTCG_2.1 and CCND2_mutTCG_2.2 primers and using DNA of the pLenti CMVTre3G eGFP Puro + ICR -CMV eGFP+1Kb PTE mut CGA construct (Cameron et al., 2018) as a template.

The second strategy was used to test all other DNA fragments and was higher throughput. To this end, we generated pLenti-Gateway vector by digesting the pLenti-ICR-Puro vector with AfeI and PmlI and recombining it with the DNA fragment containing Gateway attR cassette. This fragment was PCR amplified using primers Gateway_lenti_F and Gateway_lenti_R and DNA of the pLenti X1 Puro DEST (694-6) vector (gift from Eric Campeau & Paul Kaufman, (Campeau et al., 2009); Addgene plasmid #17297) as a template. The transgenic test constructs were produced in two steps. In the first step, DNA fragments of interest were recombined *in vitro* with pENTR1A no ccdB (w48-1) vector (gift from Eric Campeau & Paul Kaufman, (Campeau et al., 2009); Addgene plasmid #17398) digested with EcoRI using In-Fusion HD system (Clontech). The fragments of interest were PCR amplified using high-fidelity Pfu DNA polymerase (ThermoFisher Scientific), human genomic DNA as a template and sets of oligonucleotide primers indicated in Table S3. In the second step, transgenic fragments were further shuffled into the pLenti-Gateway vector via Gateway LR recombinant reaction using Gateway LR Clonase II Enzyme Mix (Invitrogen). In the case of *KCNA2* and *OVOL1*, the DNA fragments were synthesized by GeneArt (Thermo Fisher Scientific) and cloned into the pMA-RQ vector. The fragments were excised from pMA-RQ constructs using EcoRI digestion, gel purified and sub-cloned into the pENTR1A no ccdB (w48-1) vector for further shuffling into the pLenti-Gateway vector.

To produce lentiviral viral particles, 5×10^6^ 293T cells were plated in a 75-cm2 flask 24 h prior to transfection. Previously described packaging plasmids pCMV-dR8.2dvpr (6 µg) and pCMV-VSV-G (3 µg) (Cameron et al., 2018) were combined with a transgenic construct (9 µg) and co-transfected using X-tremeGene HP (Roche Applied Science) at a 1:3 ratio of DNA to the transfection reagent. After 24 h of incubation, the medium was changed. Lentiviral supernatant was collected after further 24h incubation, filtered through 0.45µm filter, and directly used for infection.

For the infection, cells were plated at a confluence of 40–60% 24 h in advance. Viral supernatants were added in serial dilutions to cells in combination with 8 µg/ml Polybrene (Millipore). Cells were subjected to single or multiple infections by adding either single lentiviral supernatant or supernatant pools (up to six different viral supernatants). After overnight incubation, the medium was changed to remove Polybrene. Transduced cells were selected for 14 days by growth in culture medium supplemented with 1µg/ml puromycin (Invitrogen).

### CRISPR/Cas9-mediated deletion of PTEs

To generate cultured cell lines lacking specific PTEs, the cells of F10 sub clone of the KBM7-1-55-S2-24 line were transfected with the Cas9/gRNA complex using electroporation by Neon Transfection system (Thermo Fisher Scientific) according to manufacture recommendations. Briefly, 2×10^5^ cells were mixed with 50pmol of Cas9 protein (IDT) and two gRNAs flanking a PTE sequence (Synthego, 25 pmol each), subjected to electric pulse (2500V for 10 milliseconds) and plated in IMDM medium without antibiotics supplemented with 15% FBS. The medium was replaced with complete medium (15% FBS, penicillin, streptomycin) after 24 hours, the cells were cultured for 2 more days and re-plated to 96-well plate after limited dilution to obtain single cell clones. DNA of the recovered clones was subjected to PCR using primers flanking the expected deletion and the nucleotide sequences of PCR products from cells homozygous for deletion alleles were determined by Sanger sequencing. Another PCR analyses was performed using primers annealing to the deleted sequence to confirm the absence of the deleted fragment anywhere in the genome. Nucleotide sequences of the gRNAs and the primers for genotyping are listed in Table S6. Genotyping results are shown in Figure S9.

### RT-qPCR, ChIP-qPCR and ChIP-seq

RT-qPCR analysis was performed as previously described in (Cameron et al., 2018). The primers used for RT-qPCR analyses are listed in Table S7.

ChIP reactions were performed as described in (Kahn et al., 2014; Schwartz et al., 2006). The antibodies used for ChIP are listed in the Table S4 and primers used for qPCR analyses are described in Table S5. MEL18 and BMI1 have nearly identical genomic binding profiles (Kahn et al., 2014; Pemberton et al., 2014). Highly specific mouse monoclonal antibodies against BMI1 protein became available from the Developmental Studies Hybridoma Bank. We therefore used this renewable and cheap reagent, instead of rabbit polyclonal antibody against MEL18, along with antibodies against CBX2 to track PRC1 binding to the transgenes.

Libraries for massively parallel sequencing (ChIP-seq) were prepared with NEBNext Ultra II FS DNA Library Prep Kit for Illumina (E7805) and NEBNext Multiplex Oligos for Illumina Index Primers Set 1 (E7335) according to manufacturer’s instructions. Briefly, 2ng of ChIP DNAs were fragmented for 20 minutes to the average length of 180bp, followed by adaptor ligation and 8 cycles of PCR amplification. Pooled libraries from MEL18 and SUZ12 immunoprecipitations with chromatins from NT2-D1 and TIG-3 cells were sequenced at NGI Sweden, SciLifeLab, Stockholm. They were sequenced from single end using 2 lines (1 flow cell) of the Illumina HiSeq2500 instrument operated in rapid 50bp read mode. Pooled libraries from all immunoprecipitations with chromatin from F10 cells and for immunoprecipitations with antibodies against RING and H3K27me3 and chromatins from NT2-D1 and TIG-3 cells were sequenced by NGI Sweden, SNP&SEQ Technology Platform in Uppsala. They were sequenced from single end using 100bp read mode of the Illumina NovaSeq 6000 instrument, v1 sequencing chemistry (Illumina) and two lanes of the SP flow cell.

### Genomic data analysis

#### Definition of bound regions

The sequencing reads were aligned to human GRCh38/hg38 reference genome using *bowtie2* (v2.2.5) (Langmead and Salzberg, 2012) set to *`--phred33 -p 8`* after which reads with ambiguous genomic positions filtered using *samtools* (v1.3.1) (Li et al., 2009) *view* command and the following parameters: *-h -b -@ 8 -q 10*. For each replicate ChIP-seq experiment, significantly enriched regions were identified with *MACS2* (v2.1.2) (Zhang et al., 2008) *callpeak* command using the following parameters: *`-f BAM -g hs --broad --min-length 1000 -B --nomodel --extsize 180 --SPMR`*. Each significantly enriched region was assigned a binding score, which corresponded to the largest sum of reads within a 1000bp window included in the region (summit window). Only regions for which the summit window was identified as significantly enriched in both replicate experiments were considered for further analyses. The “standalone” SUZ12 bound regions were defined as those at a distance of more than 100kb from any significantly enriched MEL18 region.

#### Definition of MEL18, BMI1 and RING1 binding peaks

For each ChIP-seq data set, bound regions from above were grouped together if separated no further than 100kb. ChIP-seq signals within each bound region were smoothed using *geom_smooth* function (`method = “loess”, span=0.1`) of the *ggplot2* package (https://ggplot2.tidyverse.org/). Local maxima (peaks) and local minima (valleys) in the smoothed ChIP-seq signals were identified with *stat_peaks* and *stat_valleys* functions of the same package. Each peak and valley were assigned a score representing mean ChIP-seq signal over 9 sequence positions centered on the corresponding peak or valley. The peaks with scores no lower than 50-60% of the highest peak in the same group and separated by valleys deeper than 30-35% of the scores for the flanking peaks were kept (the complete source code is presented in Supplementary script 1). This peak definition procedure returns similar number of peaks within above range of parameters. To be conservative, only peaks called at all tested combinations of parameters and identified as significantly enriched in both replicate experiments were used for further analyses. Each of these peaks was assigned a ChIP-seq signal score calculated as sum of sequencing reads within 9bp window centered on the peak position.

#### Sequence motif analyses

*STREME* version: 5.3.3 (Bailey, 2021) and the following parameters *(--verbosity 1 -- oc. --dna --p sequences.fasta --n controls.fasta --minw 4 --maxw 10 --pvt 0.05 --totallength 4000000 --time 14400*) were used to discover 4-10 nucleotide sequence motifs in DNA underneath Q4 peaks defined from the NT2 MEL18 ChIP-seq profile of the replicate experiment with the largest dynamic range. DNA sequences of 110 randomly selected 5kb genomic fragments were used as the control data set. *CentriMo* version 5.4.1 (Bailey and Machanick, 2012) executed with the following parameters (*--oc. --verbosity 1 --score 5.0 --ethresh 10.0 -- bfile sequence.fasta.bg sequence.fasta motifs.meme*) was used to calculate the number of motif matches per sequence position within 5kb DNA fragments centered on NT2-MEL18 peaks. The resulted profiles of motif matches were normalized to the total number of matches, divided in quartiles according to the corresponding peak scores, aggregated and smoothed by a rolling mean of 500 positions.

#### Calculation and plotting of the di-nucleotide frequencies

The di-nucleotide frequencies within 100bp sliding windows were calculated for 10kb DNA sequences centered on Q4 MEL18 peaks. To compare those with the genomic average, the same was done for 247 control sequences, and the latter subtracted from the frequencies calculated for DNA around Q4 MEL18 peaks. The control sequences were randomly selected from *GCF_000001405.38_GRCh38.p12_genomic.fna* as long as they did not overlap with the 10kb DNA fragments centered on the NT2-D1 MEL18 peaks (any quartile). The difference between the di-nucleotide frequencies within two independently selected sets of control sequences served as a negative control reference (right panel, Figure 7A).

#### poly(dA)/poly(dT) analysis

The frequencies of poly(dA)_5_ and poly(dT)_5_ stretches within 100bp sliding windows were calculated for 10kb DNA sequences centered on Q4-Q2 MEL18 peaks. For all instances when the putative target gene was located 5’ (upstream) of the MEL18 peak, the reverse complement sequence was used. Therefore, all target genes were oriented in the same direction, to the right of the peak. 685 control nucleotide sequences of 10kb DNA fragments not overlapping with 10kb DNA fragments centered on the NT2-D1 MEL18 peaks (any quartile) were randomly selected from *GCF_000001405.38_GRCh38.p12_genomic.fna*.

#### RNA-seq and Gene Ontology analyses

RNA sequencing data for NT2-D1 (accession # GSM3572746, (Li et al., 2019)) and TIG-3 (accession # GSM5137878, (Yano et al., 2021)) cells were downloaded from Gene Expression Omnibus (GEO) database. The sequencing reads were aligned using *bowtie2* (Langmead and Salzberg, 2012) to the *GRCh38_noalt* as index, with default arguments (*`bowtie2 -x bowtie_index/GRCh38_noalt_as −1 sequences_1.fastq.gz −2 sequences_2.fastq.gz --phred33 -q -p 10`*). *Samtools* (Li et al., 2009) was used to convert the alignments to bam format and the reads were assigned to annotated transcripts using the *featureCounts* function from the *R Bioconductor* package *Rsubread* (Liao et al., 2019), the inbuilt annotation “*hg38*” and the following parameters: *`featureCounts(outfile, annot.inbuilt = “hg38”, isPairedEnd= TRUE, nthreads=5, allowMultiOverlap = TRUE, fraction = TRUE, minMQS=20)`*. The resulted dataset was further reduced to putative PTE target genes. Those were identified as described in the Results section using gene annotation *GCF_000001405.39_GRCh38.p13_genomic.gff* table available from the National Center for Biotechnology Information (NCBI). Only annotations for which the feature column entry was either “mRNA”, “gene” or “transcript”, the “gbkey” attribute was “mRNA” or “Gene” and the “gene_biotype” attribute was “protein_coding” or missing were considered.

The Gene Ontology term enrichment was analyzed with *enrichGO* function, from *R Bioconductor* package *clusterProfiler* (Wu et al., 2021) using the following parameters: *`ont = “BP”, pvalueCutoff = 0.01, qvalueCutoff = 0.05, pAdjustMethod = “BH”, readable=TRUE`*. The gene entries from the NCBI annotation, filtered as described above, were used as a control data set. The *simplify* function was used to reduce the redundancy of the results, which were then plotted with the *cnetplot* function of the *R Bioconductor* package *enrichplot* and the following parameters: *`showCategory = 5, node_label=’all’`*

## Data access

The ChIP-seq data generated in this study are available at NCBI Gene Expression Omnibus database (GSE207401).

## ACKNOWLEDGEMENTS

We are grateful to Dr. Brent Cochran (Tufts University) for generous gift of KBM7-1-55-S2-24 cell line. We thank Dr. Jonas Sörlin (Umeå University) for help with karyotyping. Sequencing was performed by Science for Life Laboraroty at Royal Institute of Technology (KTH) and the SNP&SEQ Technology Platform in Uppsala. The facilities are part of the National Genomics Infrastructure (NGI) Sweden and supported by the Swedish Research Council and the Knut and Alice Wallenberg Foundation. This work was supported in part by grants from Cancerfonden (19 0003Pj; 22 2285 Pj), Swedish Research Council (2021-04435), Nils Erik Holmstens Forskningsstiftelsen, Basic Science-Oriented Biotechnology Research grant from the Medical Faculty, Umeå University (all to YBS) and the grant from Knut and Alice Wallenberg Foundation (2014.0018, YBS co-PI).

---

**Figure S1.**
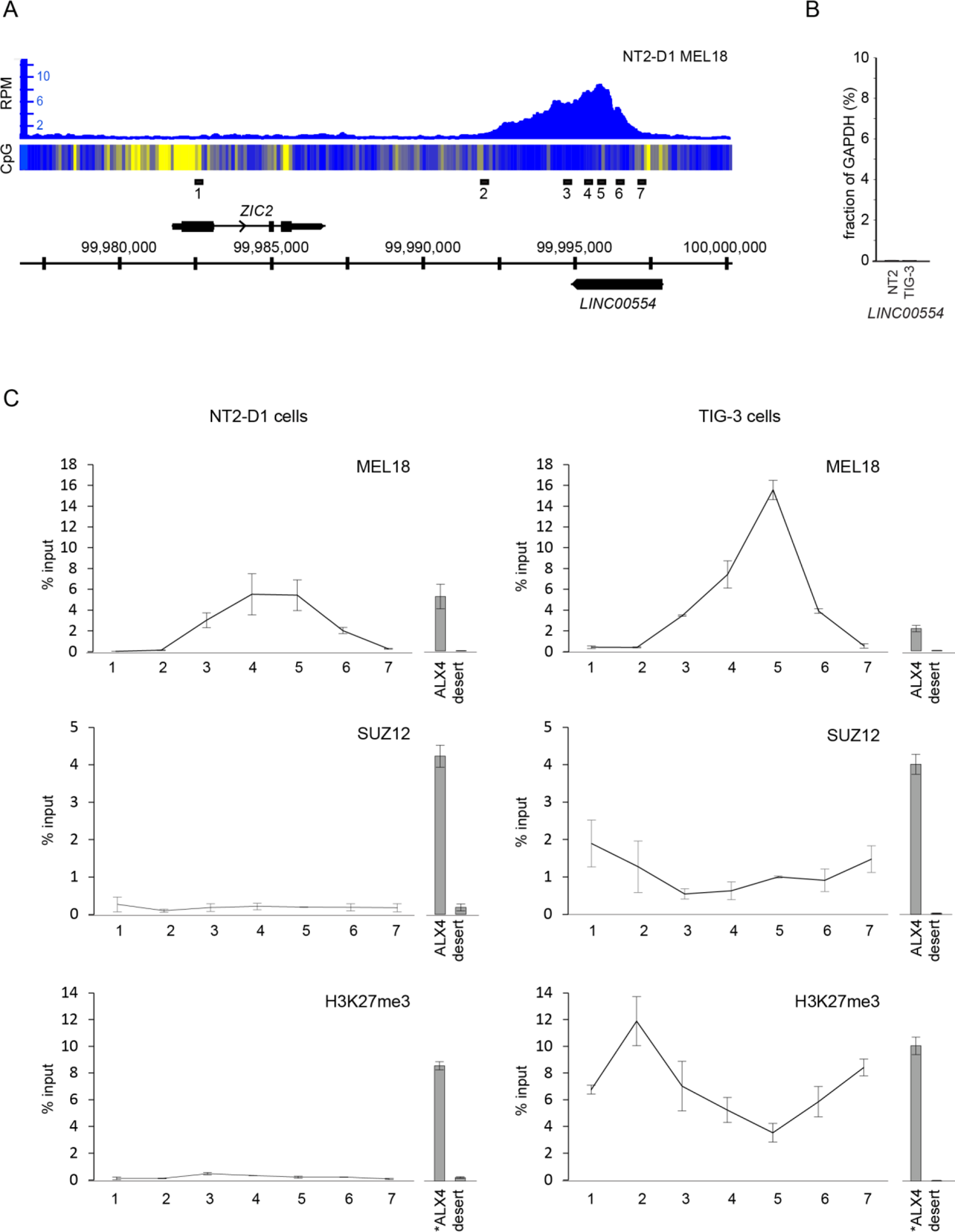
Detailed characterization of human *ZIC2* locus. **A.** ChIP-seq with chromatin from NT2-D1 cells suggests strong binding of MEL18 downstream of *ZIC2* gene. The heat-map underneath ChIP-seq profiles shows the number of CpG nucleotides within 100bp sliding window (ranging from dark blue=0 to bright yellow=15). The *ZIC2* gene is transcribed from left to right and the long non-coding RNA *LINC00554* of unknown function is transcribed from right to left. Positions of PCR amplicons (black rectangles) analyzed in **C** are shown above the scale in GRCh38/hg38 genomic coordinates. **B.** RT-qPCR measurements indicate that the long non-coding RNA *LINC00554* is not produced in either NT2-D1 (NT2) or TIG-3 cells. Values are normalized to the expression of the housekeeping *GAPDH* gene. **C.** ChIP-qPCR profiles of MEL18, SUZ12 and H3K27me3 in NT2-D1 (left column) and TIG-3 (right column) cells. The immunoprecipitation of *ALX4* gene repressed by Polycomb mechanisms in NT2-D1 and TIG-3 cells was used as positive control and Chromosome 12 gene desert was assayed as negative control. All histograms and graphs show the average and the scatter (whiskers) between two independent experiments. Note that in TIG-3 cells H3K27me3 and SUZ12 profiles are offset from the MEL18 peak into neighboring CpG-rich regions.

**Figure S2.**
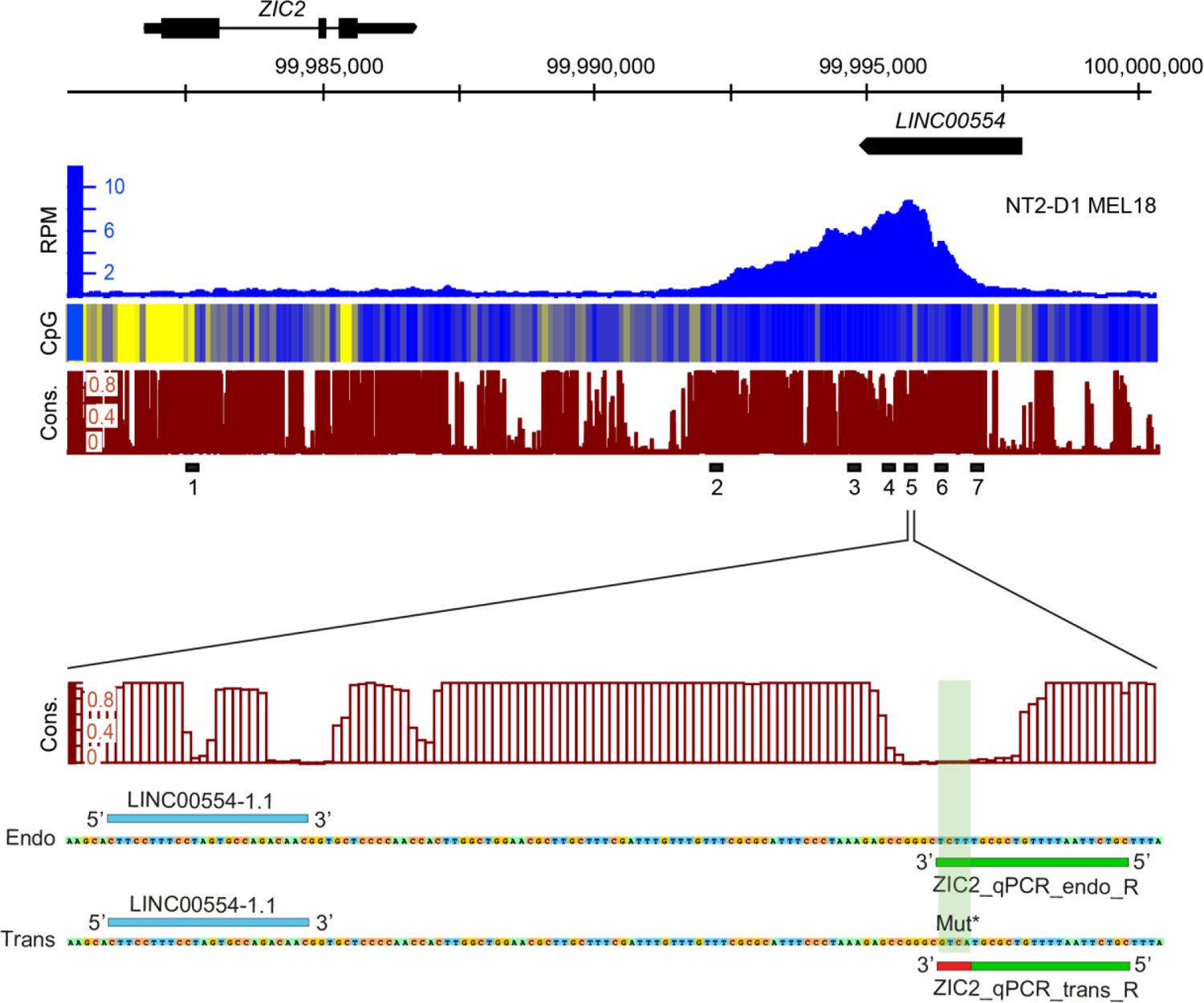
Design of the *ZIC2* transgene-specific PCR probe. In order to discriminate between the endogenous and the transgenic copy of the putative PRC1 tethering element, we used the *phastCons 100-way* conservation score to identify a small stretch of nucleotides right under the summit of the MEL18 binding peak in the *ZIC2* locus that showed little conservation within mammalian species. We posited that these non-evolutionary conserved nucleotides do not contain any important regulatory sequences and substituted them in the transgenic copy to create an annealing site for a transgene-specific PCR primer. Both the endogenous and transgenic amplicons share the same forward primer (LINC00554-1.1), but use different reverse primers (ZIC2_qPCR_endo_R and ZIC2_qPCR_trans_R respectively).

**Figure S3.**
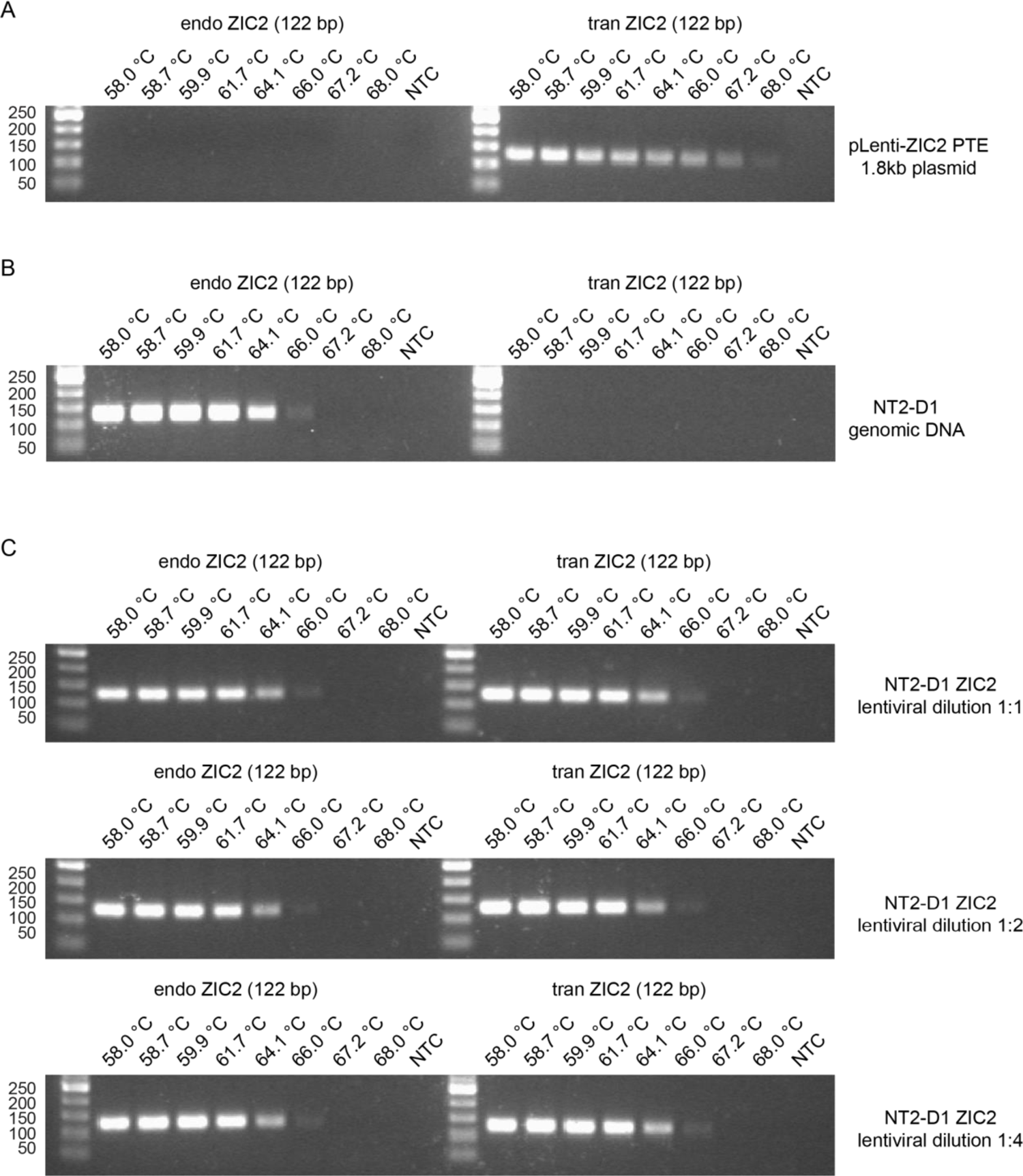
PCR genotyping of NT2-D1 cells transduced with the *ZIC2* PRC1 tethering element construct. The specificity of selected PCR primers for amplification of either endogenous or transgenic copies of the PRC1 tethering element was tested by gradient PCR using pLenti-ZIC2 PTE 1.8-kb plasmid DNA **(A)** or NT2-D1 genomic DNA **(B)** as a template. The results indicate that the PCR with transgene-specific primers amplify the product of expected size from pLenti-ZIC2 PTE 1.8-kb plasmid but not from genomic DNA. Conversely, the PCR with primer pair specific for endogenous region amplify the expected product from NT2-D1 genomic DNA but not from pLenti-ZIC2 PTE 1.8-kb plasmid. **C.** The two sets of primers were used to genotype NT2-D1 cells transduced with the lentiviral *ZIC2* PTE construct. The genotyping confirms that cells contain expected transgene. Sequences of corresponding amplicons and primers are indicated in Table S5.

**Figure S4.**
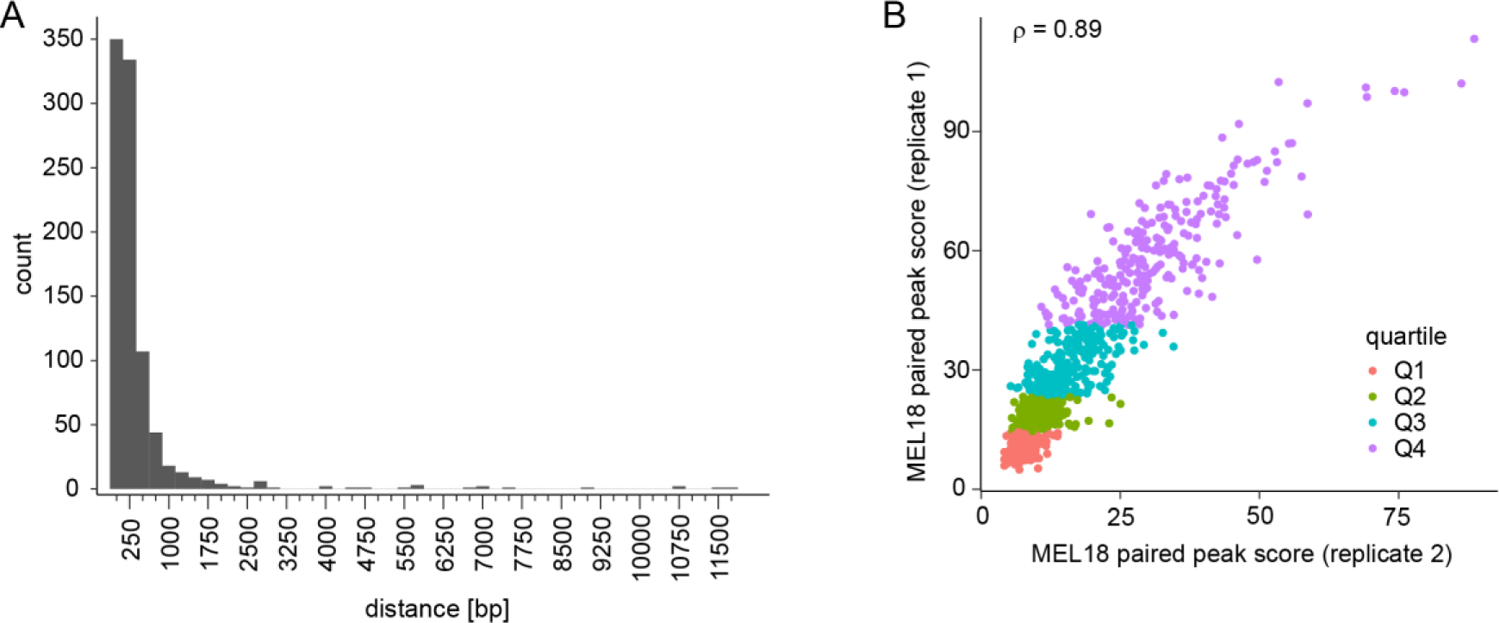
Identification of distinct MEL18 ChIP-seq peaks. **A.** Histogram of distances between the closest MEL18 ChIP-seq signal peaks from replicate experiments. **B.** The scatter plot of ChIP-seq signal scores for the paired MEL18 peaks indicates that these are highly correlated and, most often, represent the same binding site.

**Figure S5.**
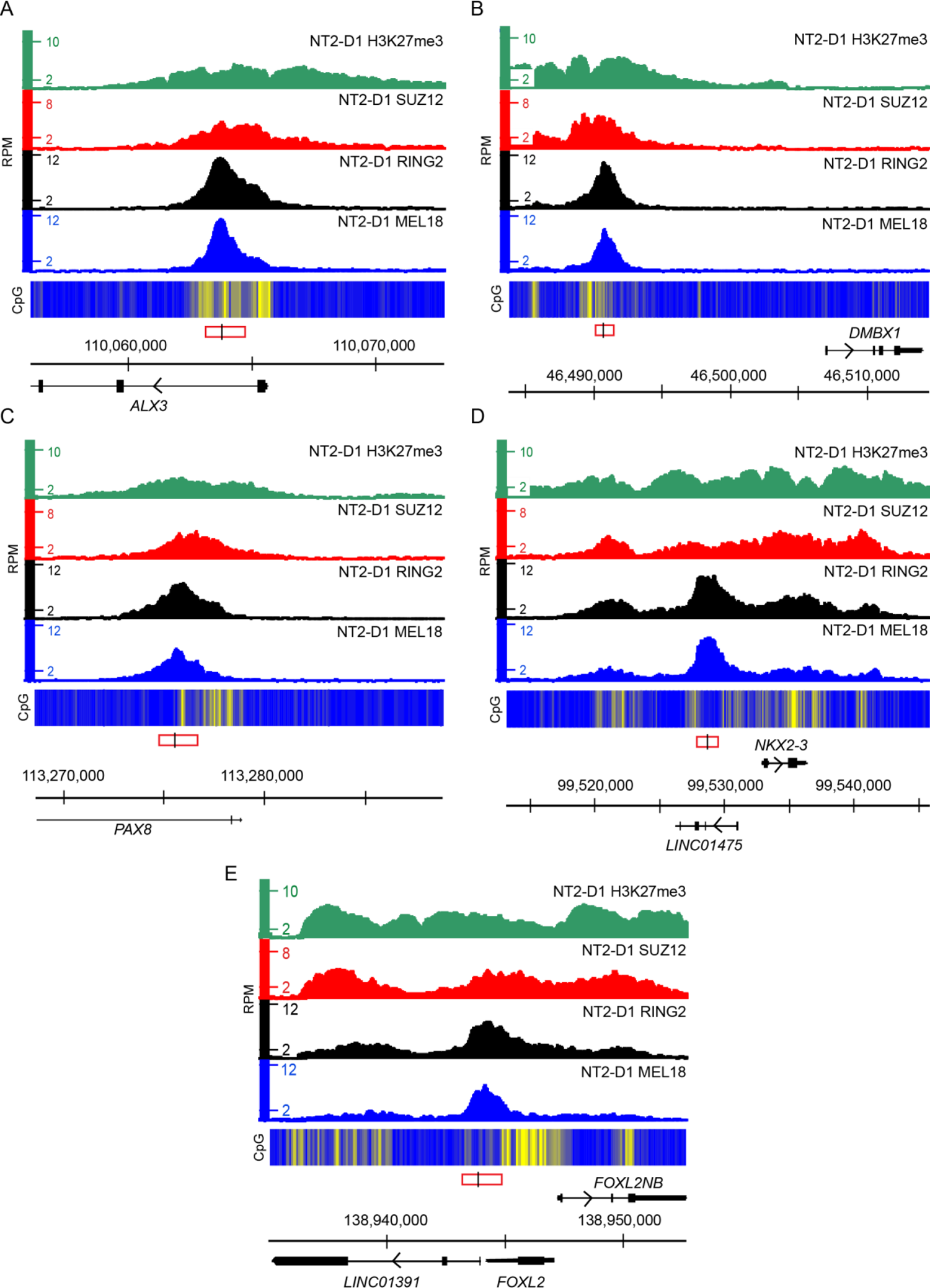
Binding of Polycomb group proteins around selected discrete Q4 MEL18 peaks. H3K27me3, SUZ12, RING2 and MEL18 ChIP-seq profiles over *ALX3* (**A)**, *DMBX1* (**B)**, *PAX8* (**C)**, *NKX2*-3 (**D)** and *FOXL2* (**E)** loci. The heat-map underneath ChIP-seq profiles shows the number of CpG nucleotides within 100bp sliding window (ranging from dark blue=0 to bright yellow=15). The regions used for transgenic assays are marked by red rectangles, with a black line indicating the position of the transgene-specific amplicon. Transcripts present in these loci are shown along the scale in GRCh38/hg38 genomic coordinates.

**Figure S6.**
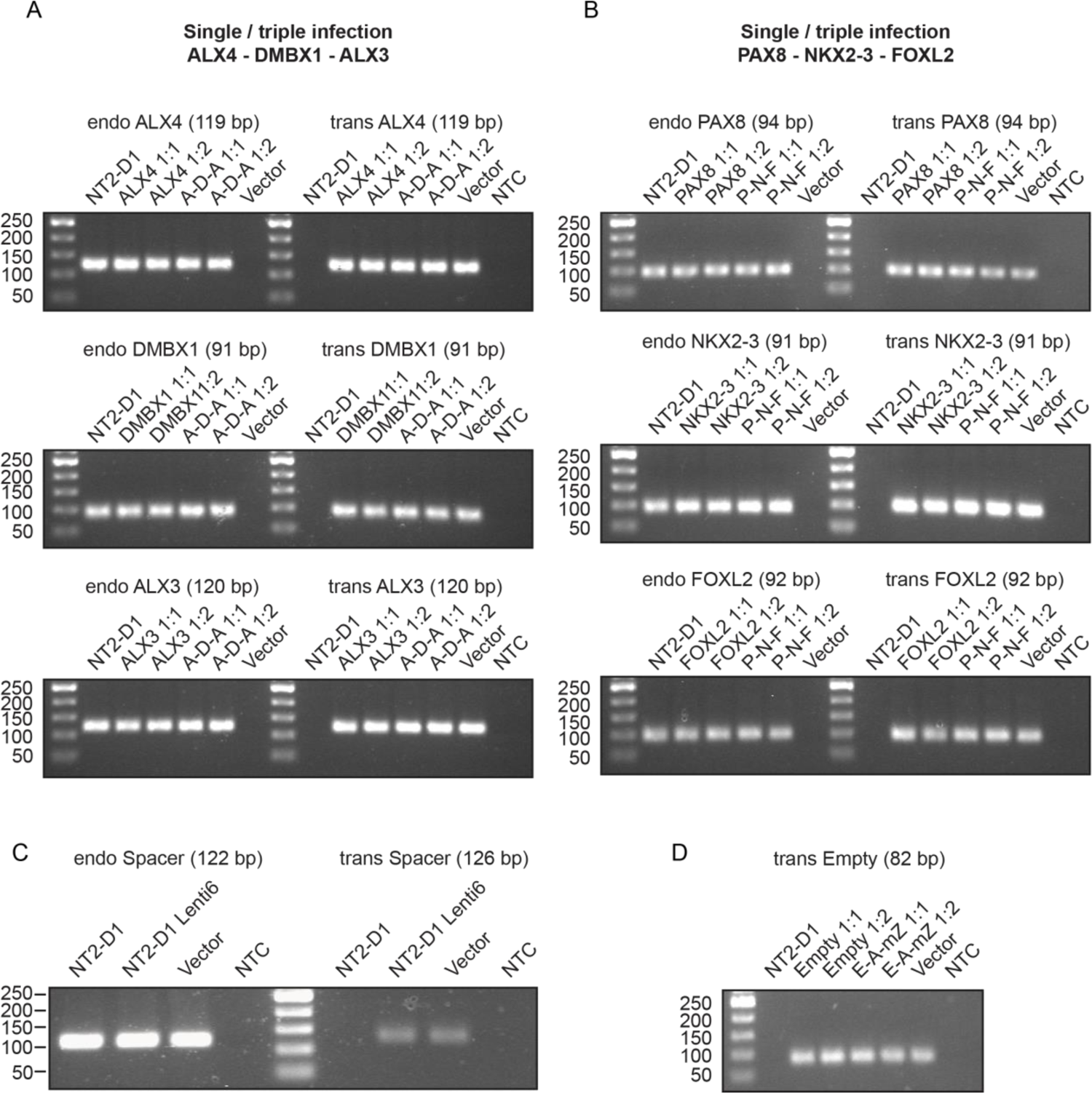
PCR genotyping of NT2-D1 cell lines transduced with Q4 PTEs. NT2-D1 cells were transduced with either a single or a pool of three different lentiviral constructs, using undiluted (1:1) or 1:2 diluted lentiviral supernatants. For all tested amplicons, NT2-D1 genomic DNA from uninfected cells was used as positive control for the endogenous amplicon (endo) and as negative control for the transgenic amplicon (trans). The correspondent lentiviral vectors were used as positive controls for the transgenic amplicons and as negative controls for the endogenous amplicons. The size of each amplicons is indicated in brackets. **A.** PCR genotyping of NT2-D1 cells transduced individually with *ALX4*, *DMBX1* or *ALX3* PTEs transgenic copies, or with the three of them together (A-D-A). **B.** PCR genotyping of NT2-D1 cells transduced individually with *PAX8*, *NKX2-3* or *FOXL2* PTEs transgenic copies, or with the three of them together (P-N-F). In both (**A**) and (**B**) the genotyping confirms the specificity of the endogenous and transgenic amplicons and that cells transduced with individual and multiple lentiviral constructs contain the expected transgenes. **C.** PCR genotyping of NT2-D1 cells transduced with lentiviral construct containing spacer DNA (NT2-D1 Lenti6). **D.** PCR genotyping of NT2-D1 cells transduced with either a single lentiviral construct containing empty vector (Empty) or a pool of three different constructs containing empty vector, *ARID3C* PTE and *ZIC2_mutTCG* PTE (E-A-mZ). Sequences of corresponding amplicons and primers are indicated in Table S5.

**Figure S7.**
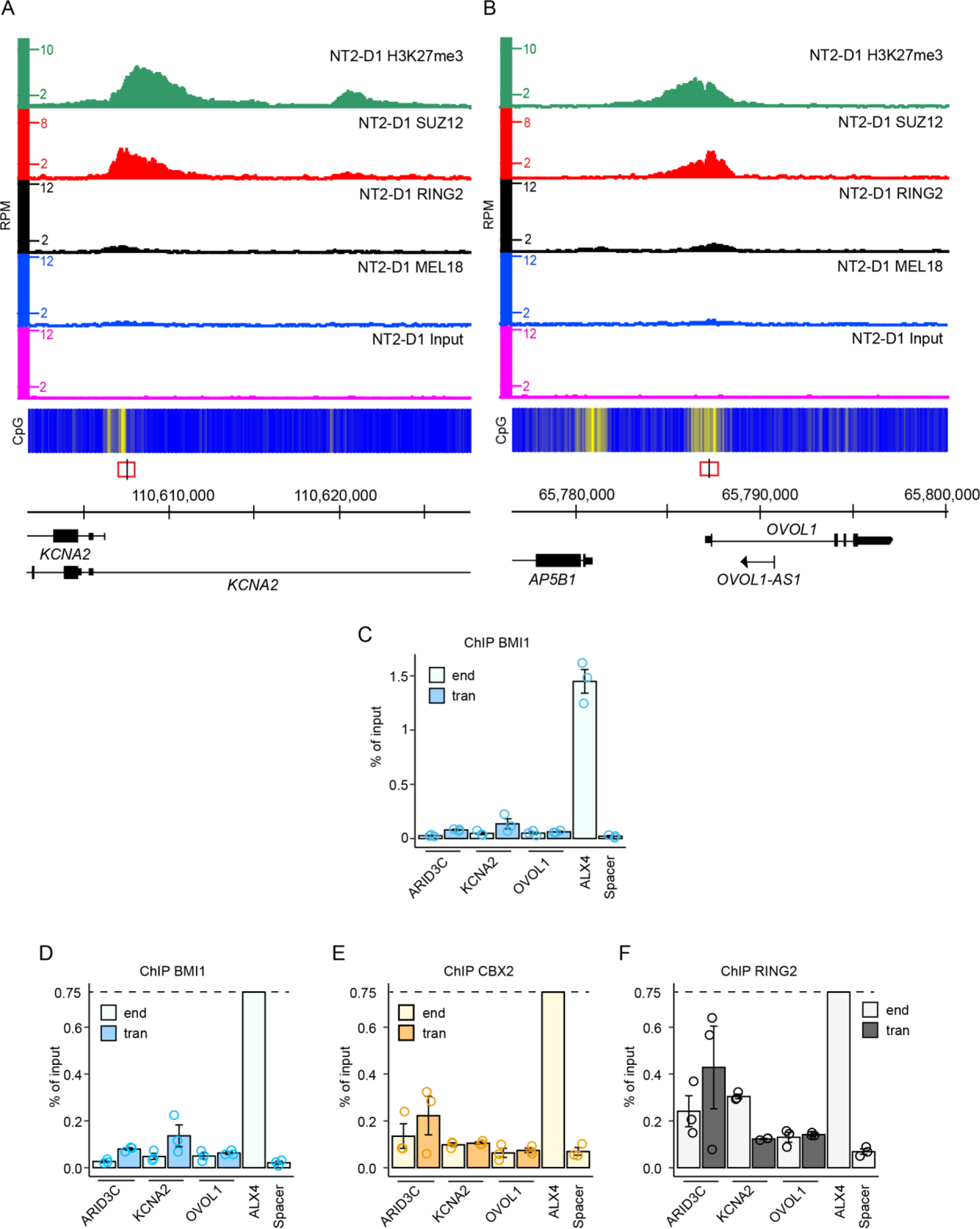
The binding of PRC2 and PRC1 to transgenic MEL18-free PRC2-bound regions. Screen-shots of the MEL18, RING2, SUZ12, H3K27me3 and Input ChIP-seq profiles around *KCNA2* (**A**) and *OVOL1* (**B**) genes in NT2-D1 cells. The fragments used for transgenic experiments are indicated by red rectangles, with genomic position of ChIP-qPCR amplicons marked by vertical black line. **C.** ChIP-qPCR experiments using BMI1 antibody as an additional canonical PRC1 component and confirming the results with CBX2 and RING2 antibodies showed in Figure 4F-G. Scaled bar-plots of yields from ChIPs with antibodies against BMI1 (**D**), CBX2 (**E**) and RING2 (**F**). Bar-plots are trimmed at y-axis value of 0.75% (indicated with dashed line).

**Figure S8.**
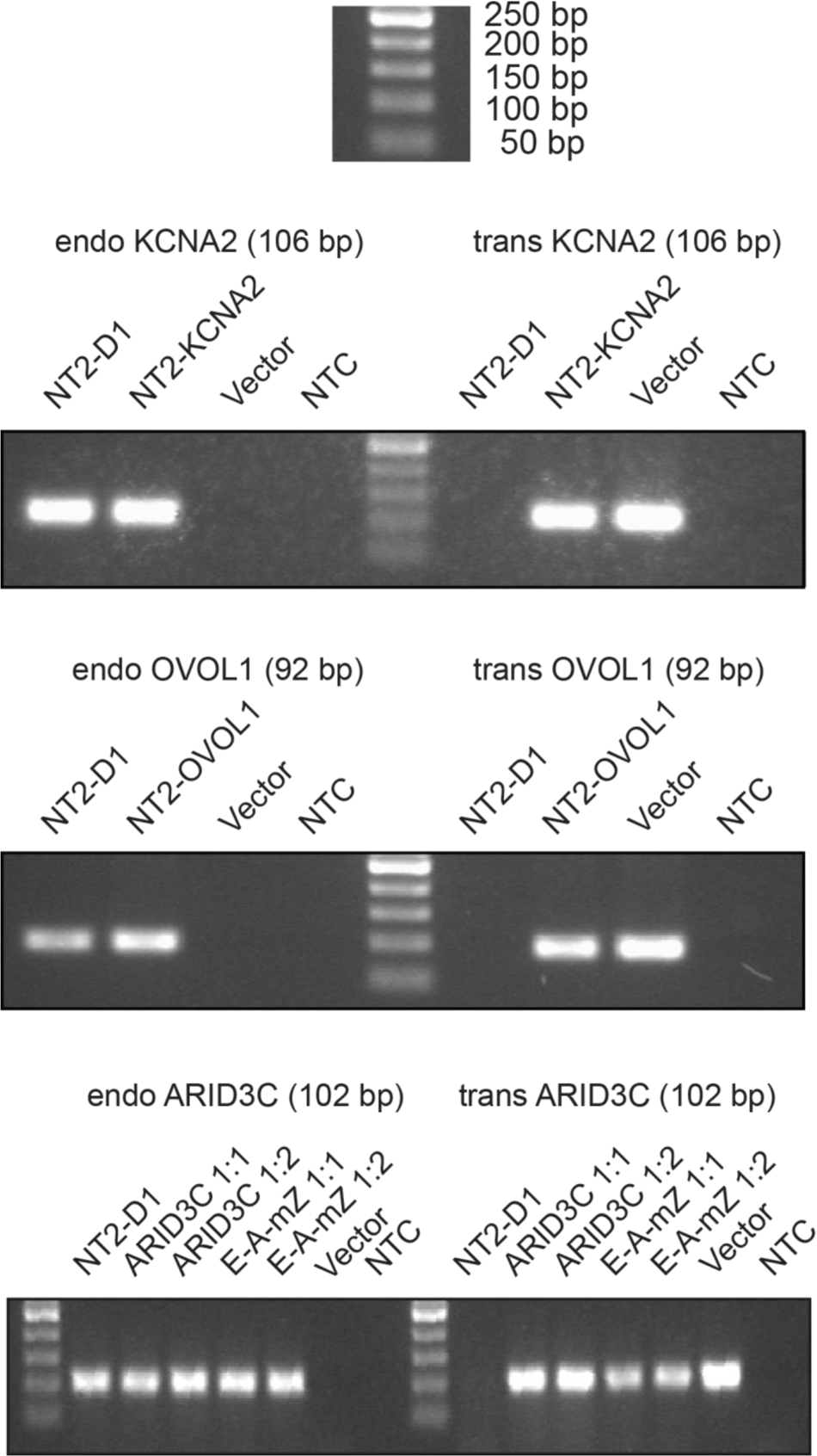
PCR genotyping of cells transduced with MEL18-free PRC2-bound regions. NT2-D1 cells were transduced with the single lentiviral construct containing *KCNA2* PTE or the single lentiviral construct containing *OVOL1* PTE or the single lentiviral construct containing *ARID3C* PTE, or the pool of three different lentiviral constructs containing empty vector, *ARID3C* PTE and *ZIC2_mutTCG* PTE (E-A-mZ). Undiluted (1:1) or two-fold diluted (1:2) lentiviral supernatants we used for some of the infections. For all tested amplicons, NT2-D1 genomic DNA from uninfected cells was used as positive control for the endogenous amplicon (endo) and as negative control for the transgenic amplicon (trans). The correspondent lentiviral vectors were used as positive controls for the transgenic amplicons and as negative controls for the endogenous amplicons. The size of each amplicons is indicated in brackets.

**Figure S9.**
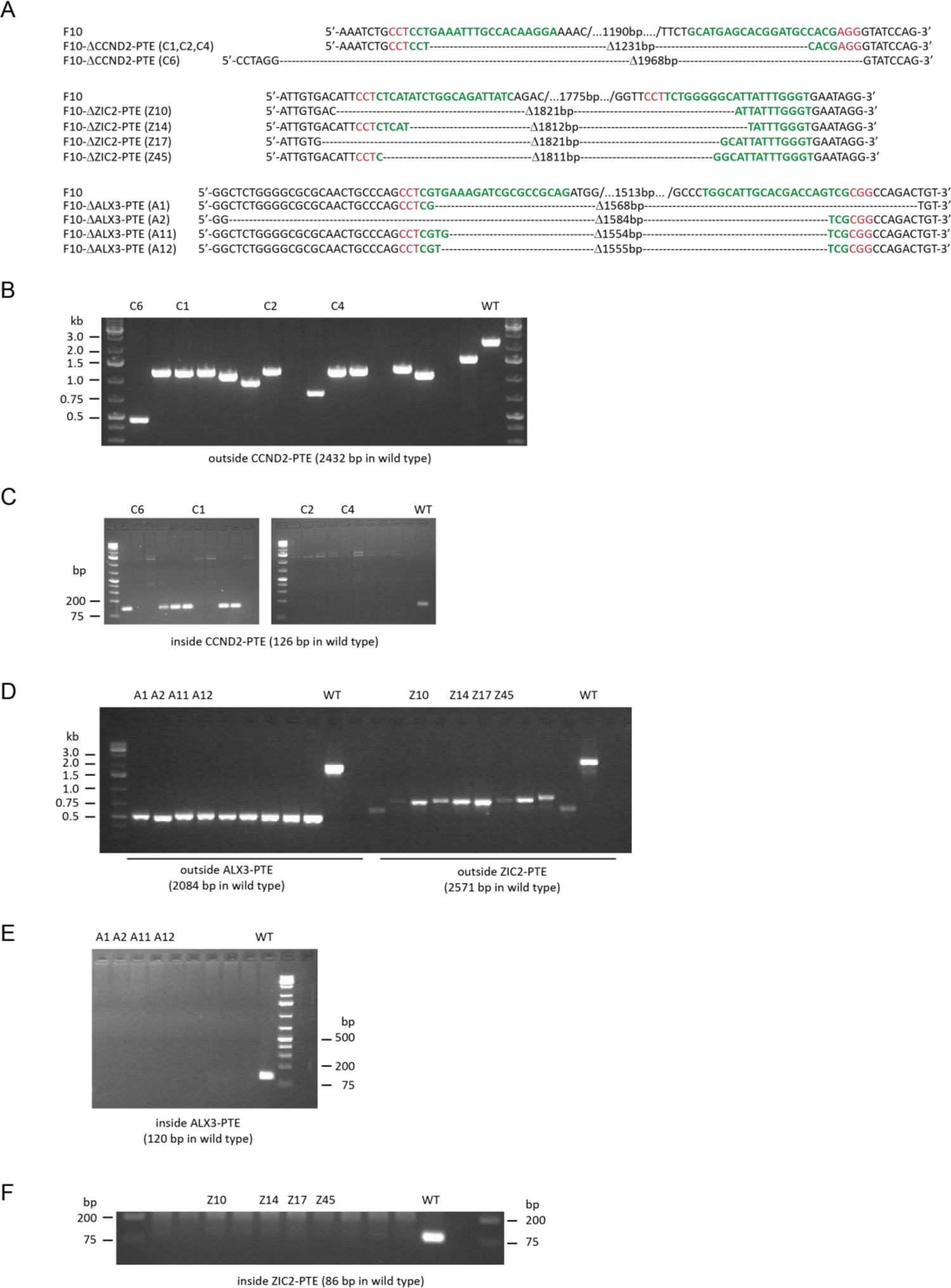
Genotyping of F10 cell lines with *CCND2*, *ZIC2* and *ALX3* PTE deletions. **A.** Nucleotide sequences of new deletion alleles used in the study. Sequences of sgRNA targets are marked in green, PAM sequences are marked in red. **B.** PCR genotyping of F10 clonal derivatives after CRISPR/Cas9-mediated genome editing around *CCND2* PTE using primers flanking the expected PTE deletion. **C.** The same analysis using primers inside the expected PTE deletion. **D.** PCR genotyping of F10 clonal derivatives after CRISPR/Cas9-mediated genome editing around *ALX3* and *ZIC2* PTEs using primers flanking the expected PTE deletions. The same analysis using primers inside the expected *ALX3* (**E**) and *ZIC2* (**F**) PTE deletions. Agarose gel lanes with products from PCRs with genomic DNA of cell lines used for further analyses are marked with corresponding clone names. Names and nucleotide sequences of the primers and sgRNAs are listed in Table S6.

**Figure S10.**
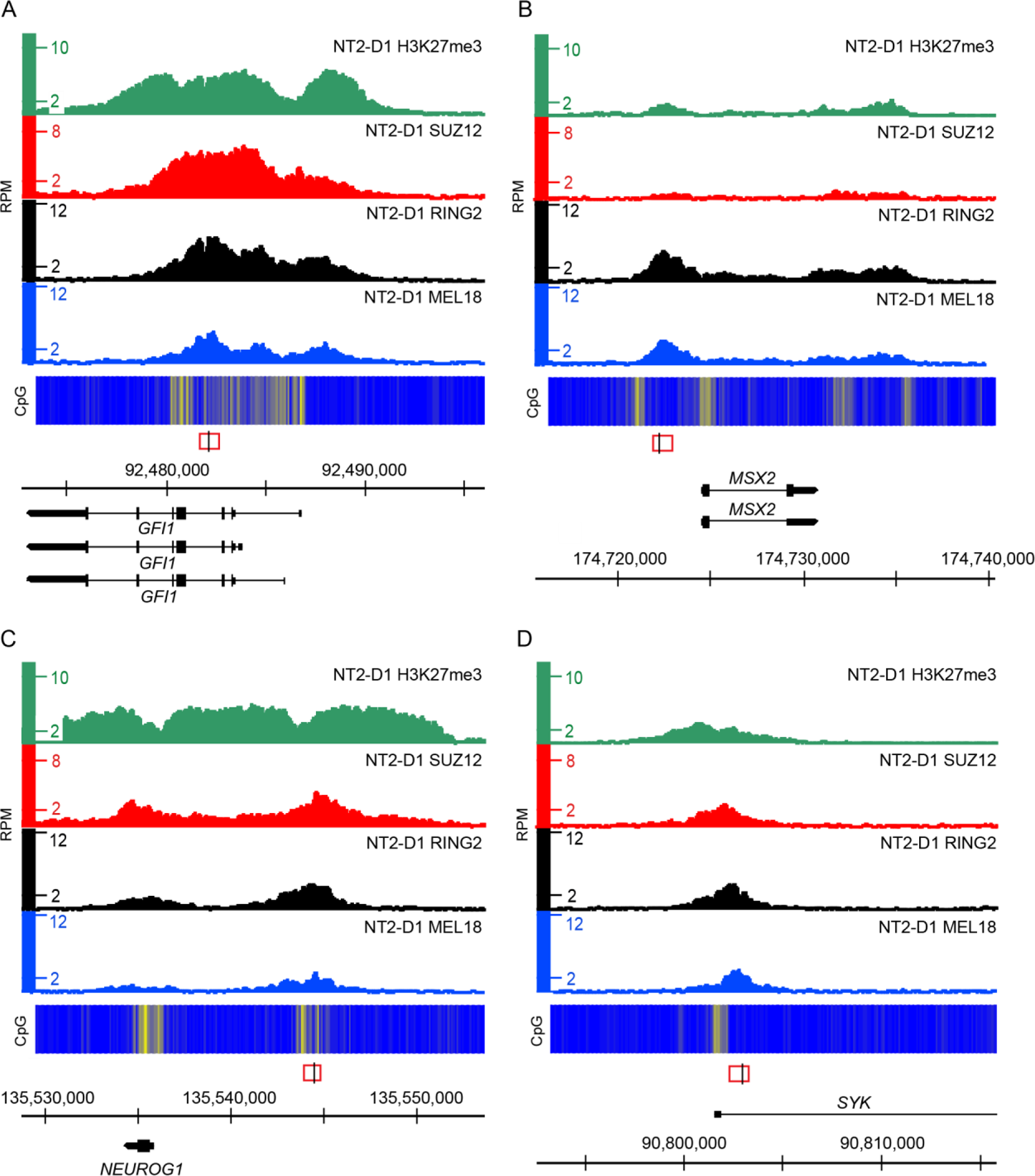
Binding of Polycomb group proteins around selected discrete Q3-Q2 MEL18 peaks. H3K27me3, SUZ12, RING2 and MEL18 ChIP-seq profiles over *GFI1* (**A)**, *MSX2* (**B)**, *NEUROG1* (**C)**, and *SYK* (**D)** loci. The heat-map underneath ChIP-seq profiles shows the number of CpG nucleotides within 100bp sliding window (ranging from dark blue=0 to bright yellow=15). The regions used for transgenic assays are marked by red rectangles, with a black line indicating the position of the transgene-specific amplicon. Transcripts present in these loci are shown along the scale in GRCh38/hg38 genomic coordinates.

**Figure S11.**
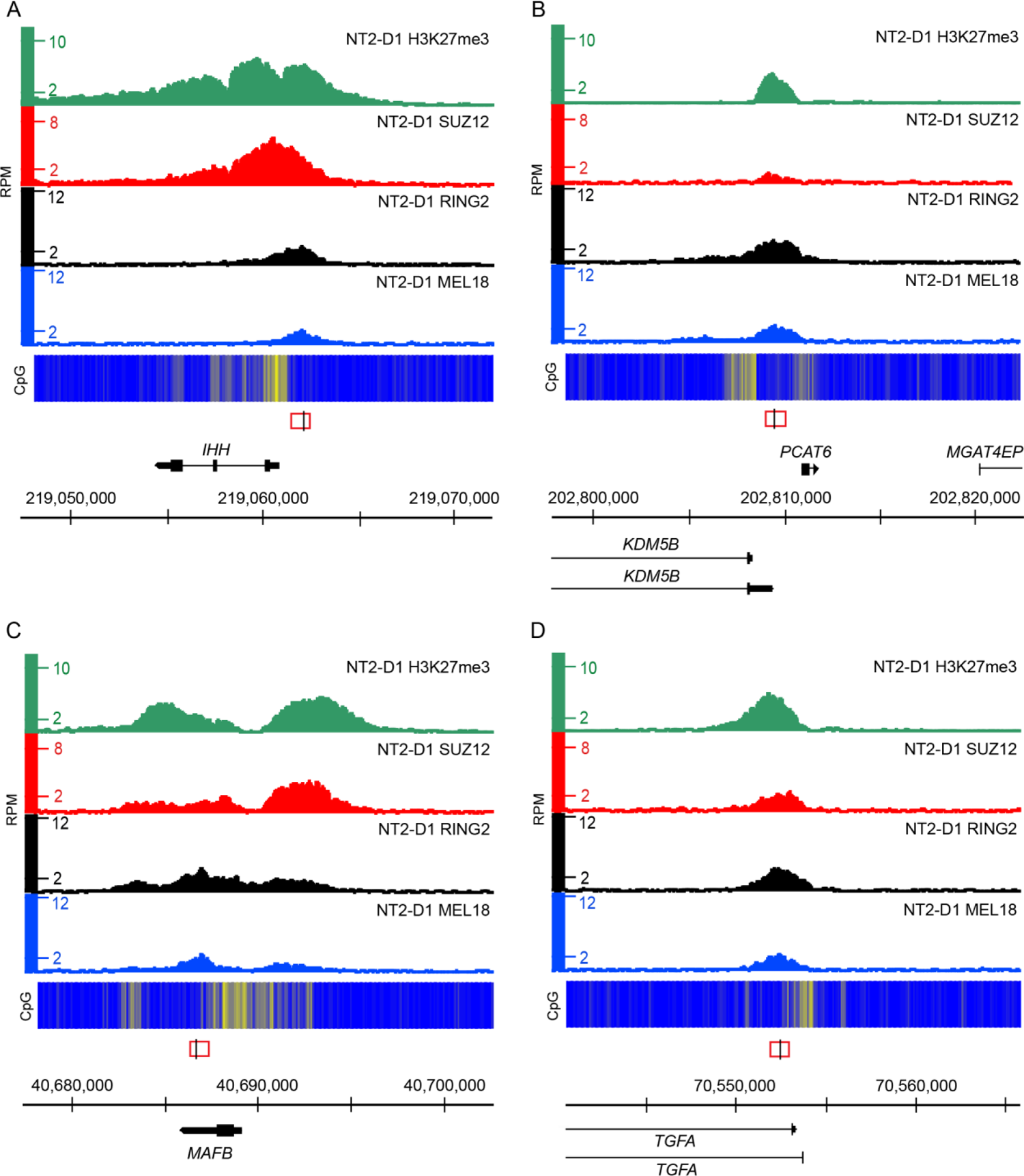
Binding of Polycomb group proteins around selected discrete Q3-Q2 MEL18 peaks. H3K27me3, SUZ12, RING2 and MEL18 ChIP-seq profiles over *IHH* (**A)**, *KDM5B* (**B)**, *MAFB* (**C)**, and *TGFA* (**D)** loci. The heat-map underneath ChIP-seq profiles shows the number of CpG nucleotides within 100bp sliding window (ranging from dark blue=0 to bright yellow=15). The regions used for transgenic assays are marked by red rectangles, with a black line indicating the position of the transgene-specific amplicon. Transcripts present in these loci are shown along the scale in GRCh38/hg38 genomic coordinates.

**Figure S12.**
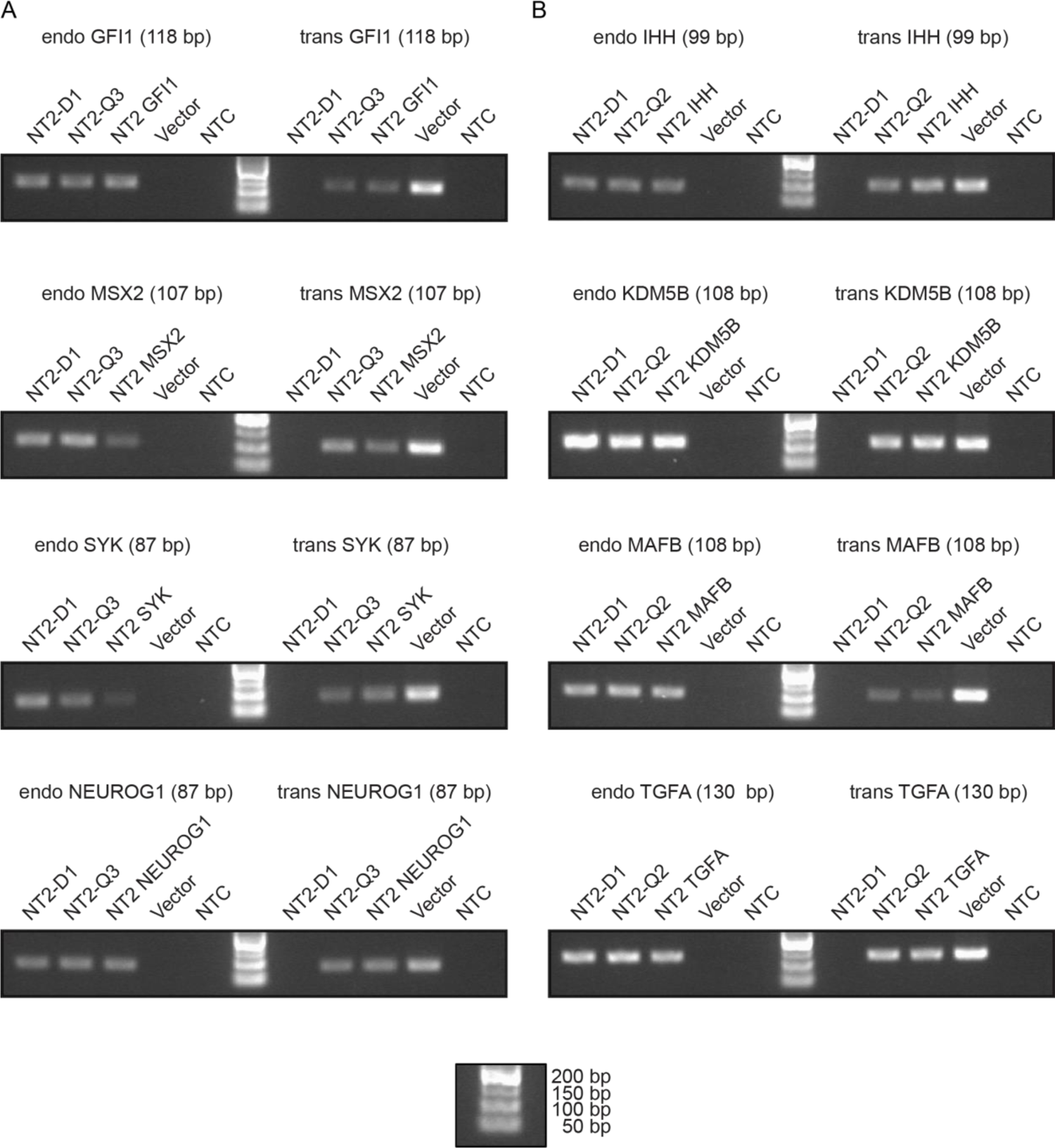
PCR genotyping of NT2-D1 cell lines transduced with Q3-Q2 PTE constructs. **A.** PCR genotyping of NT2-D1 cells transduced with *GFI1*, *MSX2*, *SYK* or *NEUROG1* PTE constructs individually and as a pool of all four constructs (NT2-Q3). **B.** PCR genotyping of NT2-D1 cells transduced with *IHH*, *KDM5B*, *MAFB* or TGFA PTE constructs individually and as a pool of all four constructs (NT2-Q2). For all tested amplicons, NT2-D1 genomic DNA from uninfected cells was used as positive control for the endogenous amplicon (endo) and as negative control for the transgenic amplicon (trans). The correspondent lentiviral vectors were used as positive controls for the transgenic amplicons and as negative controls for the endogenous amplicons. The size of each amplicon is indicated in brackets. In both (**A**) and (**B**) the genotyping confirms the specificity of the endogenous and transgenic amplicons and that cells transduced with the lentiviral constructs contain the expected transgenes. Sequences of corresponding amplicons and primers are indicated in Table S5.

**Figure S13.**
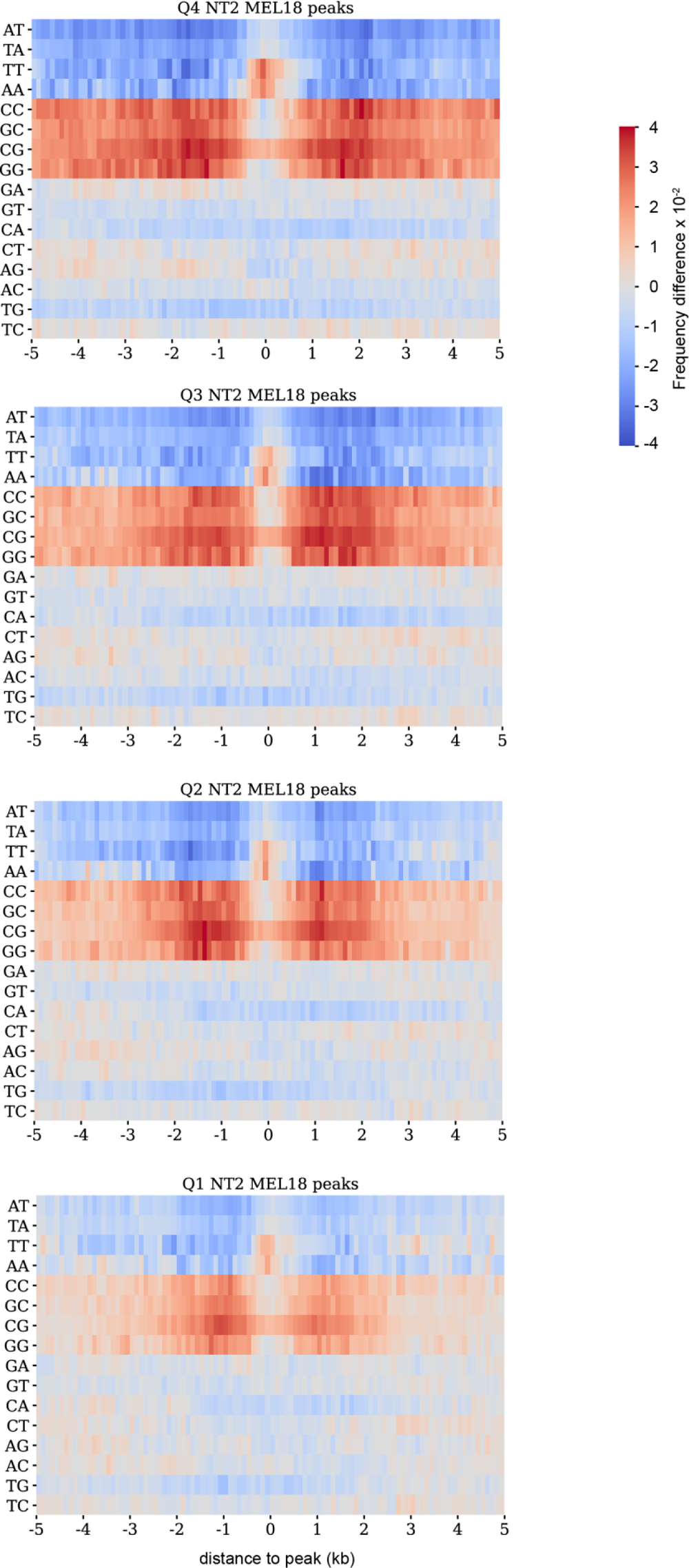
Di-nucleotide frequencies around discrete MEL18 peaks in NT2-D1 cells. Note that AA and TT enrichment under the peaks and the overall CG richness of the flanks become less prominent around less occupied (lower quartile) peaks.

**Figure S14.**
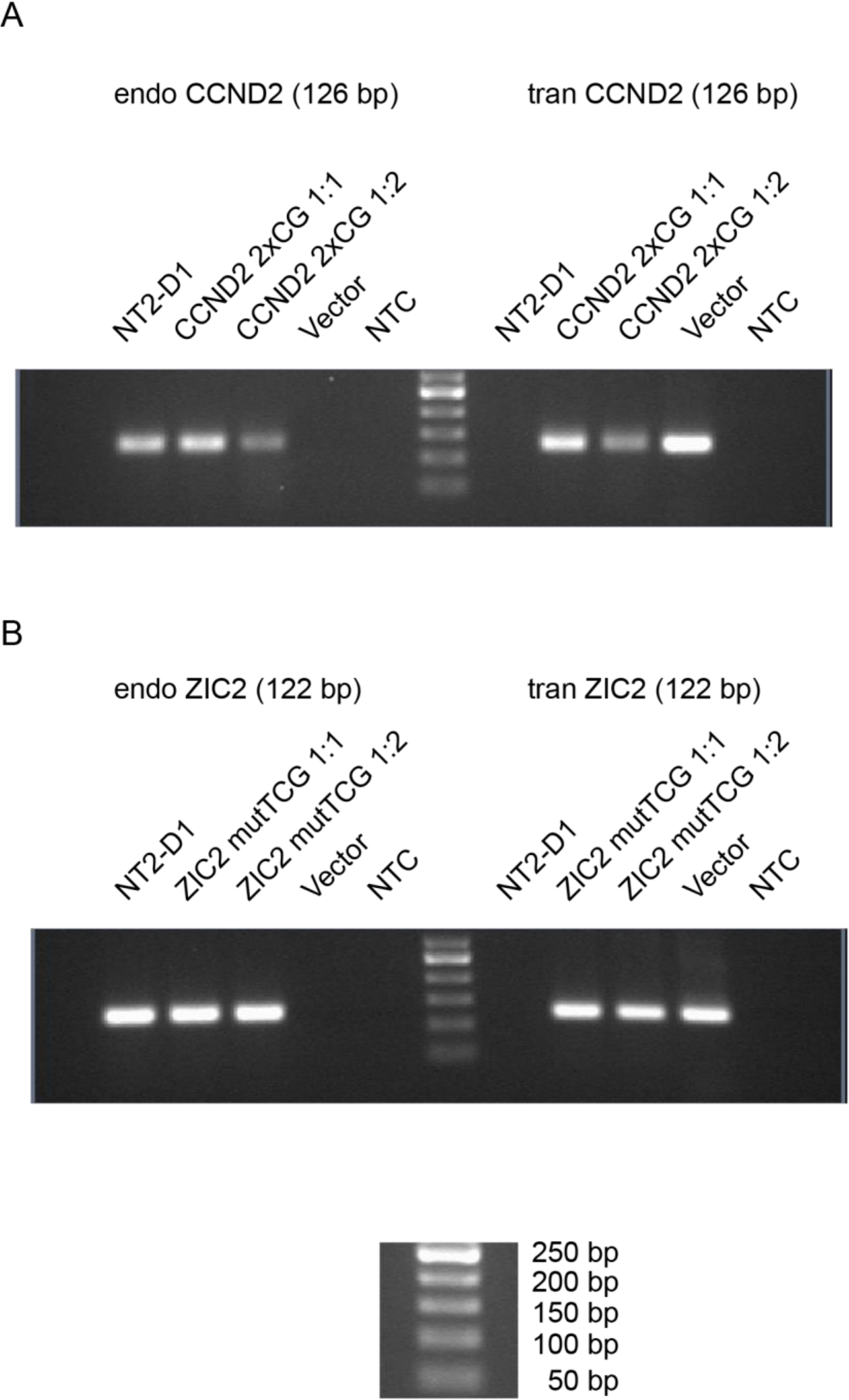
PCR genotyping of NT2-D1 cell lines transduced with mutated *CCND2* and *ZIC2* PTEs. **A.** PCR genotyping of NT2-D1 cells transduced with the CCND2_PTE_2xCGmut lentiviral construct (two best instances of “AAACGAAA” motif within *CCND2* PTE mutated) using undiluted (1:1) or two-fold diluted (1:2) lentiviral supernatants. **B.** PCR genotyping of NT2-D1 cells transduced with the pLenti-ZIC2_mutTCG lentiviral construct (the best instance of “AAACGAAA” motif within *ZIC2* PTE mutated) using undiluted (1:1) or two-fold diluted (1:2) lentiviral supernatants. For all tested amplicons, NT2-D1 genomic DNA from uninfected cells was used as positive control for the endogenous amplicon (endo) and as negative control for the transgenic amplicon (trans). The correspondent lentiviral vectors were used as positive controls for the transgenic amplicons and as negative controls for the endogenous amplicons. The size of each amplicon is indicated in brackets. In both (**A**) and (**B**) the genotyping confirms the specificity of the endogenous and transgenic amplicons and that cells transduced with the lentiviral constructs contain the expected transgenes. Sequences of corresponding amplicons and primers are indicated in Table S5.

**Figure S15.**
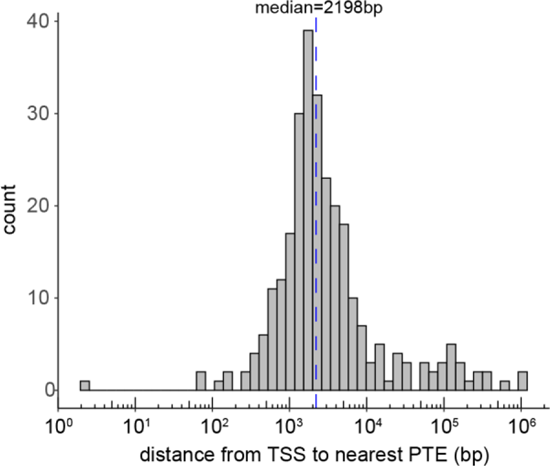
The histogram of distances between putative PTEs and TSSs of their likely target genes.

**Figure S16.**
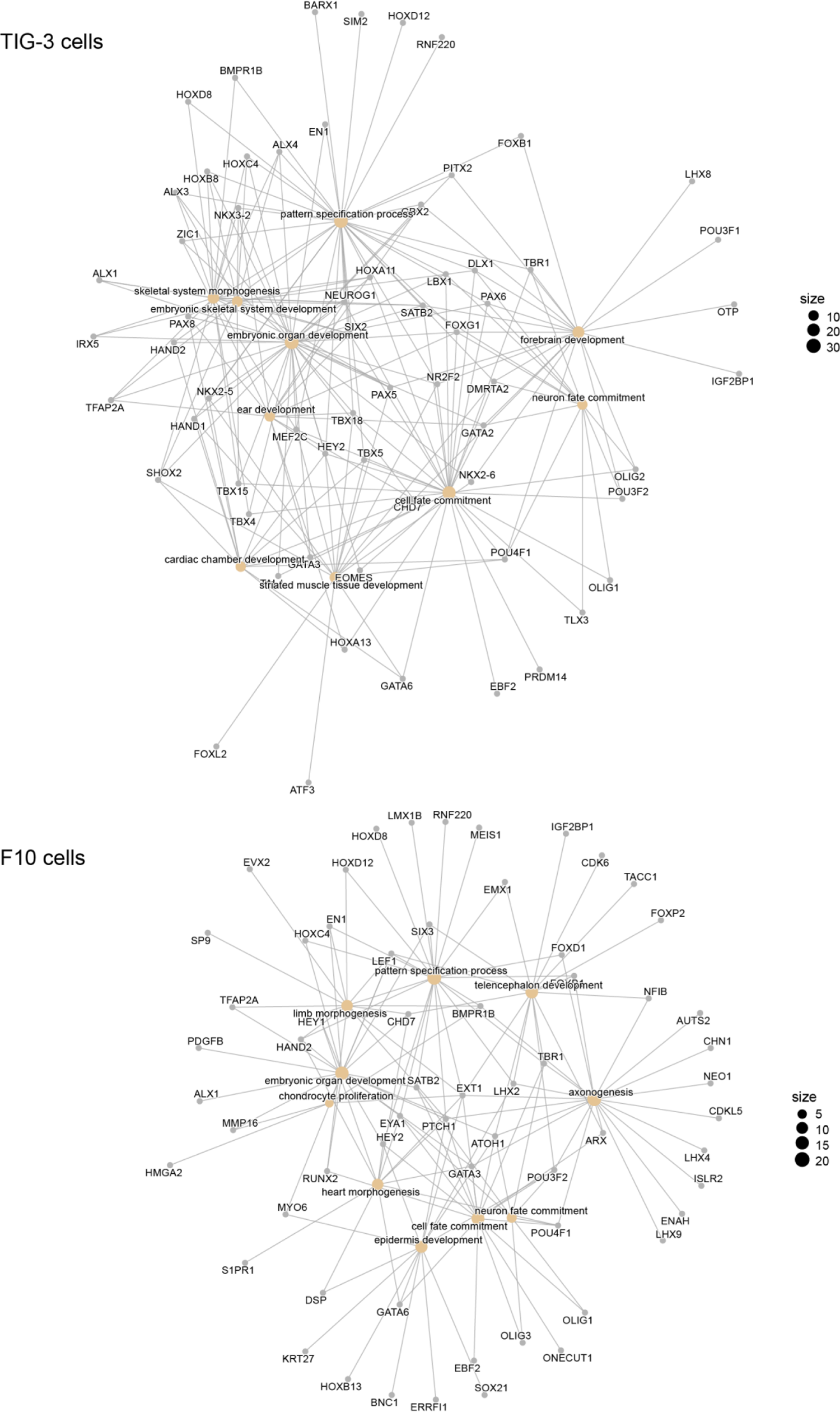
PTE equipped genes of TIG-3 and F10 cells grouped in networks by overrepresented GO terms.

## Notes

### Competing Interest Statement

The authors have declared no competing interest.

